# Temporal Interference Stimulation Can Enhance or Disrupt Human Memory Encoding as a Function of Brain Location and Frequency

**DOI:** 10.1101/2025.09.22.677714

**Authors:** Florian Missey, Eva Jouval-Missey, Mariane de Araújo e Silva, Adryelle Arantes, Claudia Lubrano, Jan Trajlinek, Samuel Klus, Ondřej Studnička, Ines R. Violante, Melanie Boly, Viktor Jirsa, Giulio Tononi, Nigel P. Pedersen, Daniel L. Drane, Adam Williamson

**Affiliations:** International Clinical Research Center, St. Anne’s University Hospital Brno, 60200 Brno, Czech Republic; School of Biomedical Engineering and Imaging Sciences, King’s College London, London, UK; Department of Neurology, University of Wisconsin, Madison, WI 53719, USA; Aix-Marseille Université, Inserm, Institut de Neurosciences des Systèmes (INS) UMR_S 1106, 13005 Marseille, France; Department of Psychiatry, University of Wisconsin-Madison, Madison, WI 53719, USA; Department of Neurology, School of Medicine and Center for Neuroscience, University of California, Davis, CA USA; Departments of Neurology and Pediatrics, Emory University School of Medicine, Atlanta, Georgia, USA; Department of Neurology, University of Washington School of Medicine, Seattle, WA USA; Center for Social and Affective Neuroscience, Department of Biomedical and Clinical Sciences, Linköping University, Linköping, Sweden

**Keywords:** Deep brain stimulation, temporal interference, bidirectional memory modulation, multi-target stimulation, Rey/Taylor figures, magnetic resonance imaging

## Abstract

Visual memory relies on synchronized interactions and rhythms between the medial temporal lobes and neocortical brain regions. Non-invasive manipulation of memory-related brain regions, specifically deeper temporal lobe regions, has been limited by the lack of precision of non-invasive neuromodulation – when targeting deeper structures, the cortex is always stimulated, never deeper structures in isolation. Temporal Interference (TI) stimulation, a novel non-invasive technique, uses high-frequency carrier fields to deliver targeted, physiologically relevant neuromodulation via amplitude-modulated envelopes at specific brain regions. Here, we investigate TI’s impact on figure memory encoding in 70 healthy participants using the Rey-Osterrieth and Taylor Complex Figure tasks, with TI applied in several brain regions independently and simultaneously – allowing investigation of combinations of medial temporal lobe and neocortical brain regions. Interestingly, higher frequency TI envelopes (130 Hz offset) targeting bilateral hippocampi and temporal cortices significantly impair recall (p = 6.54e-04), while lower frequency TI envelopes (5 Hz offset) targeting only the bilateral hippocampi significantly enhance recall (p = 0.0447). Stimulation using other combinations of medial temporal lobe and neocortical regions showed no effect, underscoring the critical role of frequency and focality of non-invasive brain stimulation and correct target selection. Finally, functional MRI reveals strong differences between the effects of 130 Hz and 5 Hz envelopes, specifically in hippocampal BOLD signals, brain connectivity, default mode, and attentional networks. These findings demonstrate TI’s ability to bidirectionally modulate memory encoding through precise frequency and target tuning, offering a powerful tool for cognitive neuroscience and potential therapeutic applications for memory disorders.

**Graphical Abstract:**
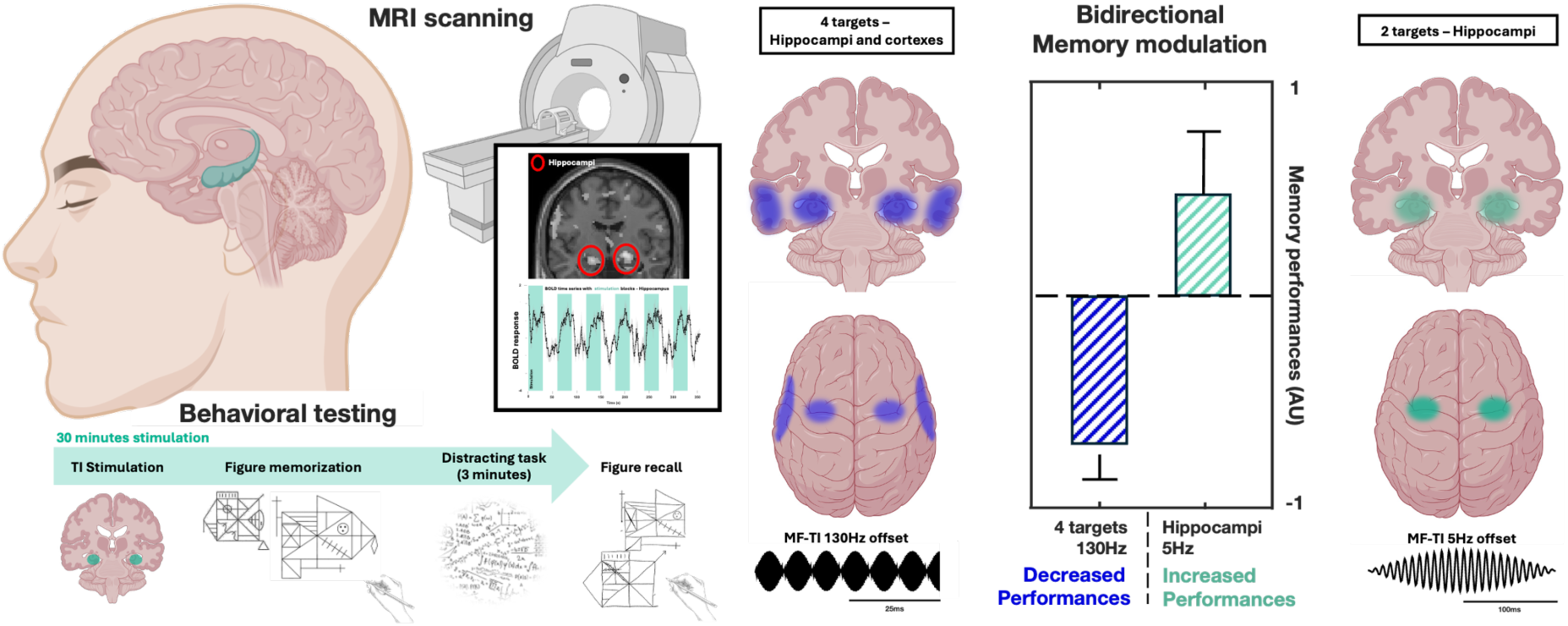
Temporal Interference stimulation for bidirectional memory modulation assessed via behavioral testing and MRI scanning.

## 1. Introduction

Memory relies on synchronized activity across brain regions, including the hippocampus, perirhinal cortex, parahippocampal cortex, and retrosplenial cortex, which form a network critical for episodic memory, navigation, and planning (Ranganath & Ritchey, 2012; Vann et al., 2009). The hippocampus serves as a central hub, orchestrating memory encoding and consolidation through interactions with neocortical areas and subcortical structures like the amygdala and anterior cingulate cortex (Preston & Eichenbaum, 2013; Squire & Wixted, 2011). Disruptions to these networks, as seen in Alzheimer’s disease or following bilateral medial temporal lobe damage (e.g., patient H.M.), lead to profound memory impairments, while enhanced connectivity can improve performance, highlighting the importance of precise, targeted interventions (Scoville & Milner, 1957; Dresler et al., 2017).

Brain stimulation offers a powerful approach to modulate memory by influencing neural plasticity, but traditional methods face limitations: invasive techniques like deep brain stimulation lack flexibility, while non-invasive methods, such as transcranial direct current stimulation, often lack depth and precision (Kuo & Nitsche, 2012; Suthana et al., 2012). The timing, target, and frequency of stimulation critically determine its effects, with evidence suggesting that hippocampal stimulation can impair memory if mistimed, whereas theta-frequency stimulation during encoding can enhance it (Jacobs et al., 2016; Kirov et al., 2009). These findings underscore the need for a non-invasive method capable of targeting deep brain structures with high specificity and frequency control to bidirectionally modulate memory processes.

In this study, we employ Temporal Interference (TI) stimulation, a non-invasive method which uses high-frequency carrier fields to deliver focal neuromodulation to deep brain regions (Grossman et al., 2017; Missey et al., 2021). We specifically designed a multi-focal TI approach that allowed us to simultaneously target different regions of the memory network within the medial temporal lobes and neocortical brain regions.

This multifocal design revealed frequency-specific effects with bidirectional modulation of memory encoding. Using the Rey-Osterrieth and Taylor Complex Figure tasks, we show that low-frequency TI (5 Hz offset) targeting the hippocampi enhances recall, while high-frequency TI (130 Hz offset) targeting both hippocampi and temporal cortices impairs it, highlighting the critical role of frequency and target selection.

Functional MRI further reveals that TI induces frequency-dependent hippocampal BOLD signal changes and alters connectivity in default mode and attentional networks. These findings demonstrate TI’s capacity to precisely tune neural dynamics, enhancing or impairing memory encoding based on stimulation parameters. This approach offers a powerful tool to probe memory networks and holds potential for therapeutic interventions in cognitive disorders.

## 2. Material and Methods

### Behavior experiments

#### Participants

70 participants were recruited from St Anne University Hospital students/staff and Masaryk University students in accordance with the approved protocol EKV-2022-096 by Masaryk University Research Ethics Committee. Before taking part in the study, participants were screened for memory issues, neurological and psychiatric conditions, and substance abuse. The study followed the ethical guidelines set by Saint-Anne University Hospital (SAUH) and adhered to the principles of the 1964 Helsinki Declaration and its subsequent revisions. Informed consent was secured from the participants before any experimental procedures.

#### Stimulation parameters

Multi-focal TI was performed using eight DS5 devices (Digitimer®, UK) driven by function generators (FG, Keysight®, USA), one FG generates two waveforms and runs two DS5s at the same time. Each pair of DS5s was allocated to a different target in the brain. Stimulation electrodes were selected based on Finite Element Modelling (FEM) generated in Sim4Life software (see below) and within the 256-sensors Geodesic Sensor Net (MagStim EGI) (Fig. 1A).

**Figure 1:**
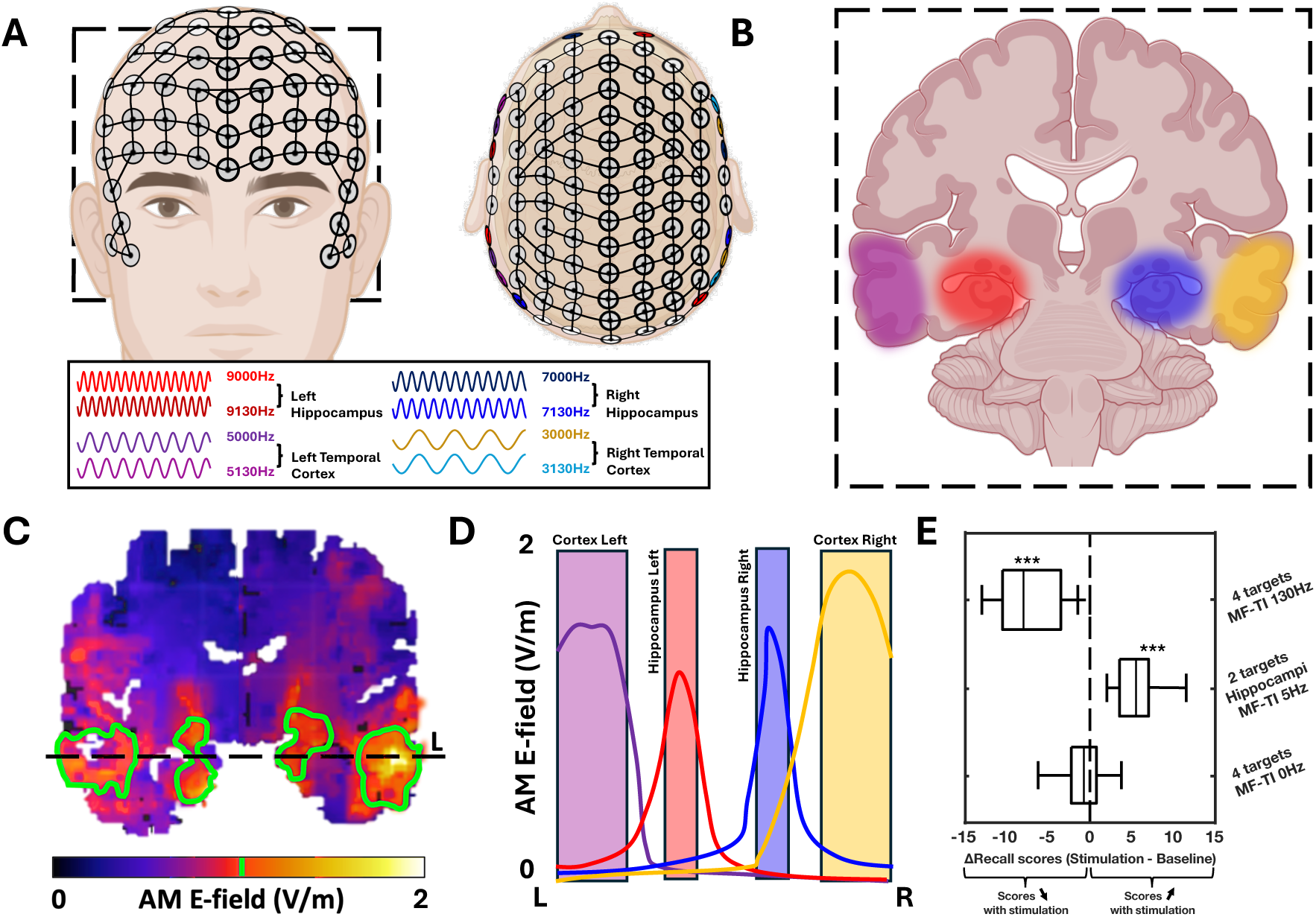
Multi-focal Temporal Interference stimulation design. Multi-focal Temporal Interference was delivered using a high-density EEG cap where specific electrodes were allocated for stimulation. The stimulation frequencies were selected with a 2 kHz offset to avoid unwanted cross-stimulation between the different targets (**A**). Our multi-focal TI protocol enabled the targeting of 4 different brain areas simultaneously (**B**). Before the experiments, finite-element modeling was performed to assess the stimulation in the different targets (green outline)(**C**). The Amplitude-Modulated (AM) field is plotted across regions on the horizontal axis, highlighting the multifocality of the approach (**D**). Multi-target TI stimulation can efficiently modulate memory by decreasing or increasing performance based on the offset frequency used (**E**).

##### High-frequency four-target stimulation

TI stimulation frequencies were centered on 3000, 5000, 7000, and 9000 Hz with a 130 Hz offset to modulate four different targets (Fig. 1B and C). As high frequencies are known to penetrate biological tissues more due to decreased impedance (Lempka et al., 2009), we selected 9000 and 7000 Hz carriers to target bilateral hippocampi and, 3000 and 5000 Hz for cortical TI stimulation (Fig. 1). Stimulation maximal amplitudes were set at +/- 2mA and adjusted so that participants had no stimulation-related cutaneous sensations. The stimulation session started 30 minutes before the Rey/Taylor task was administered and was slowly ramped until reaching +/- 2mA. Stimulation was stopped right before the figure recall.

##### High-frequency two-target stimulation

TI stimulation frequencies used were 7000/7130 Hz, and 9000/9130 Hz to modulate two different targets, either bilateral hippocampi or bilateral temporal cortices. The stimulation amplitudes and durations remain the same as for the four-target protocol.

##### Low-frequency four-target stimulation

Low-frequency TI stimulation involved the same carrier frequencies as for the high-frequency four-target stimulation, *i.e.,* centered around 3000, 5000, 7000, and 9000 Hz, except with an offset frequency at 5 Hz. This frequency is within the human theta rhythm range and was described as facilitating encoding by Violante et al., 2023. Bilateral hippocampi and temporal cortices were targeted. The frequency distribution, maximum amplitude, and stimulation duration were similar as presented above.

##### Low-frequency two-target stimulation

This was the same as high-frequency two-target stimulation, however with an offset frequency of 5 Hz. The frequency distribution, maximum amplitude, and stimulation duration were similar as presented above.

##### Sham stimulation

Sham stimulation uses the same carrier frequencies, 3000, 5000, 7000, and 9000 Hz but with no offset. The stimulation protocol was otherwise the same as those described above.

##### Protocol

Participants were randomly assigned, in a double-blinded manned, to receive multi-focal TI stimulation at either a low amplitude modulation frequency (Δf=5 Hz), high amplitude modulation frequency (Δf=130 Hz), or sham with no offset (Δf=0 Hz) – 10 participants were assigned per group for a total of 7 groups and 70 participants.

#### Behavioral testing

The Rey-Osterrieth and Taylor Complex figure task protocols were designed to evaluate visual memory encoding, retention, and recall abilities. These figures were chosen for their intricate designs, requiring considerable cognitive effort to encode accurately, and have shown no differences in complexity (Hubley et Jassal, 2006; Zhang et al., 2021). These standardised neuropsychological instruments, are among the most widely used, have demonstrated robust psychometric properties across diverse populations and are extensively validated in clinical applications, making them useful for measuring the potential therapeutic efficacy of novel memory interventions. Before starting, the participants were given specific instructions on how the experiment would proceed and what was expected from them throughout the entire task. Out of the 70 participants, none of them knew the figures in advance and participants only took place in the study once.

##### Complex figure memory task

The task began with the complex figure copying and encoding. Participants were randomized to be shown either the Rey-Osterrieth figure or Taylor Complex figure and given as much time as they needed to copy the complex geometrical patterns. This was followed by a distracting task to prevent working memory effects. The distracting task lasted for three minutes and consisted of mathematical operations. Subsequently, participants were required to draw the figure they copied earlier.(Fig. 2A and 3A). The quality of the reproduced figure provides quantitative and qualitative data on the participant’s memory performance based on objective feature scoring that captures aspects of both overall form and details.

**Figure 2:**
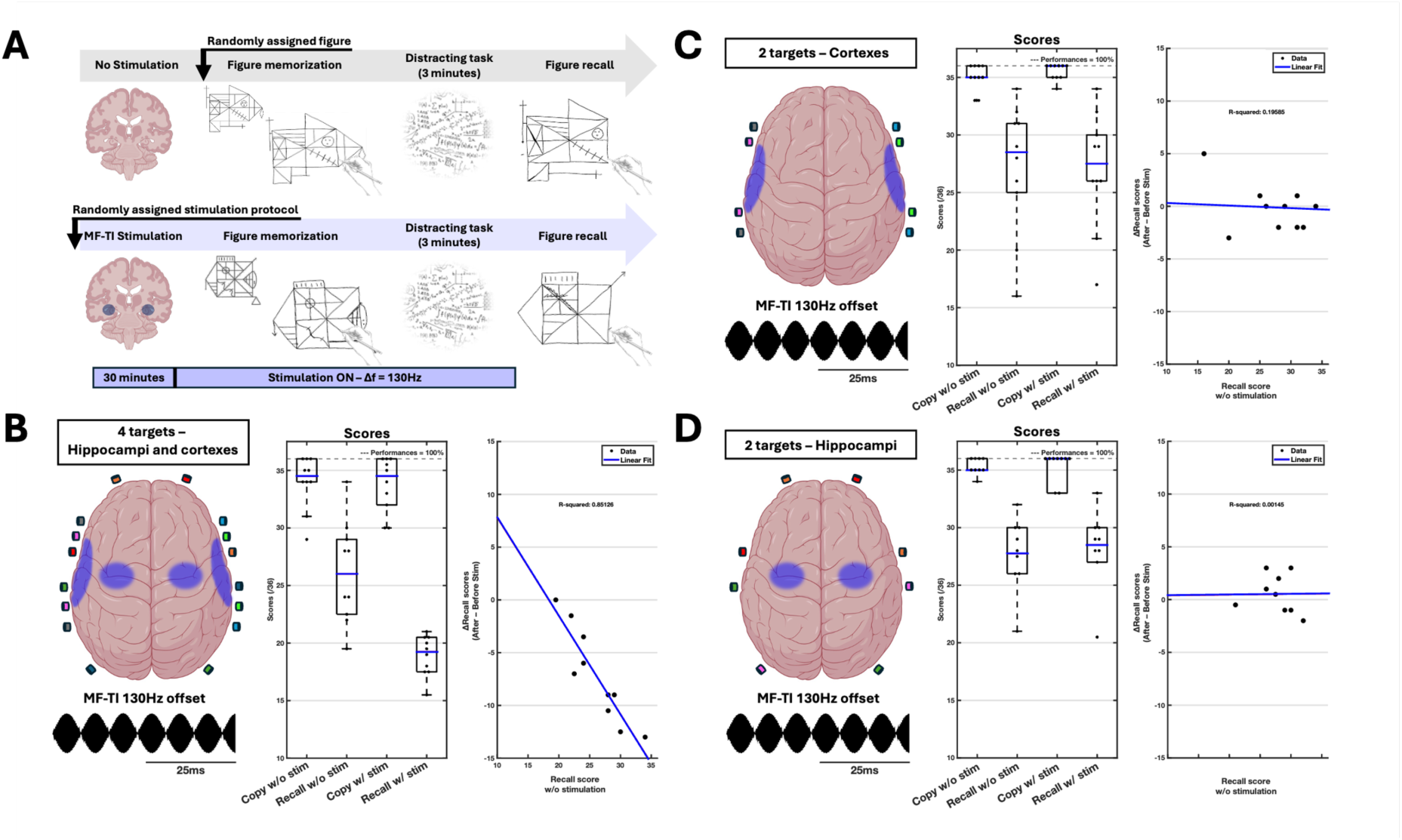
Study protocol and Rey/Taylor figure scores analysis after high-frequency multi-focal TI stimulation. Multi-focal TI stimulation was provided in 4 distinct targets, bilateral hippocampi and bilateral temporal cortices. The offset used for high-frequency stimulation was 130Hz (**A**). The baseline and TI experiment scores were gathered and compared without pre-processing or via linear regression of the recall scores before and after stimulation. The raw scores and the correlation between the delta in recall scores and the initial recall during the baseline experiment has been calculated for the 4 targets, the 2 target cortices, and the 2 target hippocampalprotocols (**B, C,** and **D** respectively). As can be seen in C and D, 130 Hz envelope, a frequency we have previously used in therapeutic applications (Parkinson’s disease, Epilepsy) showed no diminished ability to recall when focally targeting either cortex of hippocampus. However, as seen in B, when all four regions are targeted, clear deficits in recall were observed during the stimulation condition.

**Figure 3:**
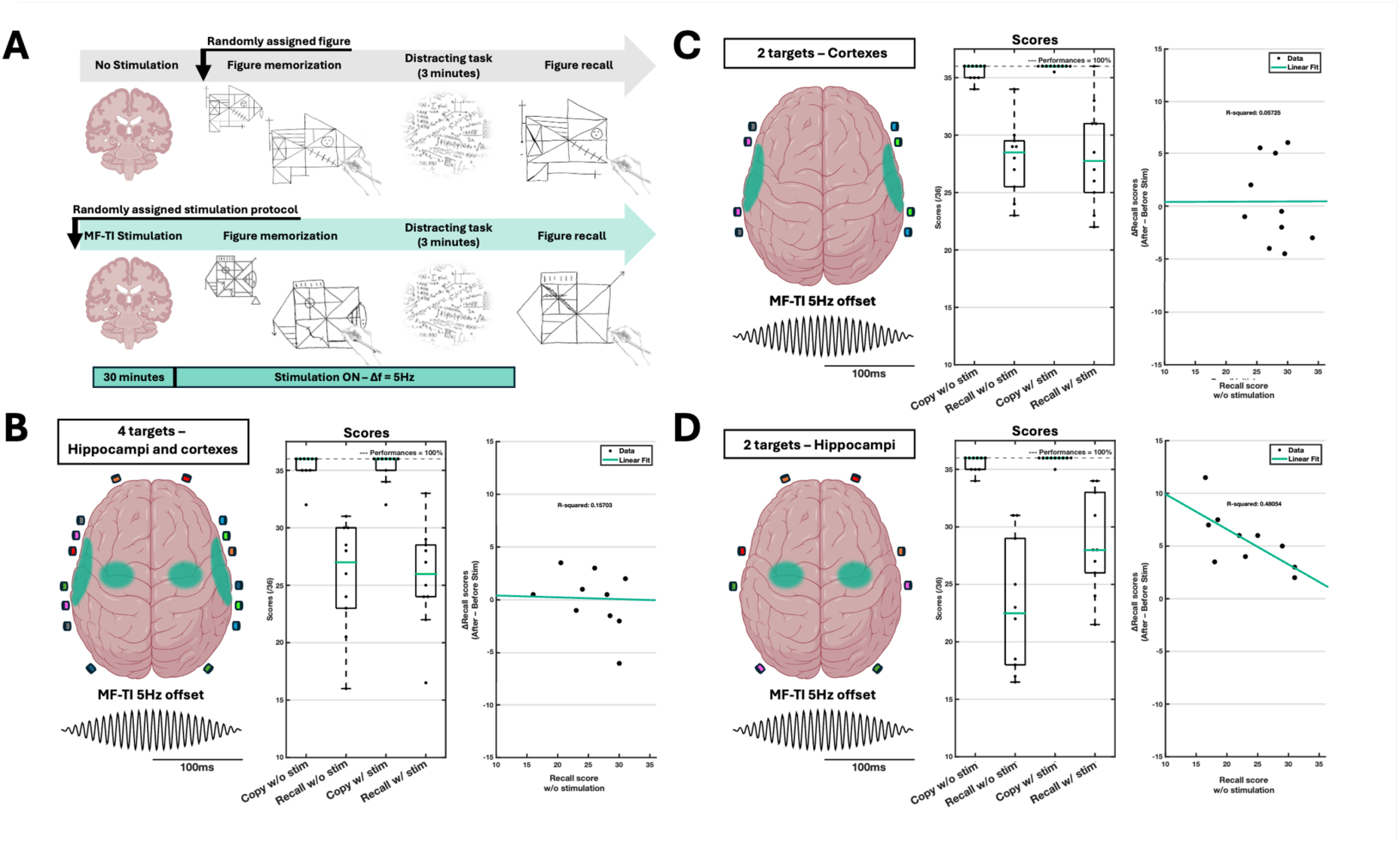
Study protocol and Rey/Taylor figure scores for multi-focal TI low-frequency stimulation. Multi-focal TI stimulation was provided in 3 distinct targets, bilateral temporal cortices, bilateral hippocampi and bilateral temporal cortices and hippocampi. The offset used for low-frequency stimulation was 5 Hz (**A**). The baseline and TI experiment scores were gathered and compared without pre-processing or via linear regression of the recall scores before and after stimulation. for the three targeting setups (**B, C,** and **D**). As can be seen in B and C, when targeting cortex or hippocampus and cortex, no enhanced recall was observed, the increase is seen only in D, where neuromodulation is applied focally to the hippocampus where enhanced recall is observed.

##### Behavioral task design and conditions

In the baseline condition, no stimulation was introduced, and the task protocol took place as mentioned above (Fig. 2A and 3A – top panel) – the baseline condition always occurred before the TI condition. In the TI conditions, a 30-minute stimulation of one of the five multi-focal TI stimulation patterns presented in the stimulation parameters section (High-Frequency, Low-Frequency, or Sham x 2-targets or 4-targets) was introduced. The stimulation started 30 minutes before the memorization/study phase and was stopped right before the recall/test phase, thus preferentially targeting encoding (Fig. 2A and 3A – bottom panel). Participants were randomly assigned to one of the two figure for the baseline condition and got the remaining one for the TI condition, so that participants did not show practice effects by figure repetition. Participants attended the two sessions: baseline and TI stimulation. Thus, this experimental design was developed to test the following specific hypotheses regarding the effects of multi-targeted temporal interference brain stimulation on the encoding of novel visual stimuli and its downstream impact on short-term recall: (i) low-frequency TI stimulation (5 Hz offset) targeting only the bilateral hippocampi should enhance recall performance compared to baseline, based on prior evidence that theta-range frequencies facilitate memory encoding (Violante et al., 2023); (ii) high-frequency TI stimulation (130 Hz offset) should lower brain activity/connectivity, impairing recall performance compared to baseline, consistent with findings that higher frequencies can disrupt synchronized neural activity in memory networks (Jacobs et al., 2016); and (iii) sham stimulation should have no significant effect on recall, underscoring the importance of frequency specificity. These hypotheses were tested by comparing recall performance between the baseline (no stimulation) and experimental (TI stimulation) conditions.

##### Outcome measures

3-minute recall in the Rey-Osterrieth Complex Figure (Rey figure) and Taylor Complex Figure (Taylor figure) involves evaluating an individual’s ability to reproduce a previously copied complex figure from memory after a specified delay period. The scoring is based on comparing the recalled reproduction to a standardized template, focusing on accuracy in detail placement, spatial arrangement, and overall completeness. Copy and 3-minute recall were scored quantitatively, on a scale ranging from 0 to 36, where higher scores indicate better accuracy and completeness in reproducing the figure with the template or from memory after a delay. These scores are interpreted in comparison to established normative values, providing insights into participants’ performances in learning and recall. The Complex Figures were scored by a neuropsychologist using a double-blind design (i.e., neither the participants nor the neuropsychologist was aware of the stimulation parameters) and following standard test scoring criteria. (Fig. S1 and S2).

#### Statistical analysis

The study included 70 participants for the behavioral task (mean age = 27.93 years, SD = 7.57; 35 women). All participants were right-handed and had no known history of neurological or psychiatric disorders. Baseline characteristics, including age, sex, education level, and baseline performance on the Rey/Taylor task, showed no significant differences between the experimental conditions (p > 0.05). All participants safely tolerated the stimulation at +/- 2mA without any sensation on the skin or adverse effect post-stimulation; in no case the current had to be lowered or the stimulation stopped during the experiment.

The baseline and TI conditions were compared for our three stimulation protocols on 3-minute recall scores. Before any statistics, a descriptive analysis was run for each condition, including mean, median, standard deviation, and range calculation. Subsequently, all group data were tested using Shapiro-Wilk tests to evaluate the normality of the immediate 3-minute recall score distributions within the conditions. Depending on the output of the normality tests, parametric tests (t-tests) or non-parametric (Wilcoxon-Mann-Whitney) tests were conducted to compare the scores between the conditions. A further subgroup analysis was performed to examine potential covariates of the observed effect, such as age and baseline cognitive performance. To account for multiple comparisons across the different stimulation conditions (baseline or TI stimulation *x* TI protocol), we applied the Bonferroni correction to control for type I error, and all displayed p-values are corrected.

#### Finite element simulations

Finite element simulations of multi-focal temporal interference were performed on Sim4Life software (Zurich MedTech AG), using an electro-ohmic quasi-static solver, which solves the equation ∇σ∇ϕ = 0, where σ is the local electrical conductivity, and the E-field is obtained as E=-∇ϕ. As ohmic currents dominate current displacements, Maxwell’s equations were approximated with σ≫ωɛ (ω : angular frequency), with σ≠0 as the E-fields are only determined in the lossy domain. The human model used for the simulation is part of the Virtual Population library from Sim4Life (Jeduk V4.0). As part of the ITI’S Foundation database of tissue properties (Hasgall et al., 2011), conductivities and permittivities of all biological tissues were automatically assigned to the model. Stimulation electrodes were modeled based on the size of electrodes on the 256-sensors Geodesic Sensor Net (MagStim EGI). Electrode selection (based on the 256 electrode net) was first approximated to focalize an amplitude-modulation Temporal Interference stimulation in the target areas. Positions were then improved depending on the output of the approximate simulation until an optimized placement was found (Fig. 1D and E). Total-current-normalized data were obtained from the stimulation, which was carried out at active electrodes under

Dirichlet boundary conditions. **Equation 1** from Grossman & al., 2017 was used to get the maximum envelope modulation amplitude.

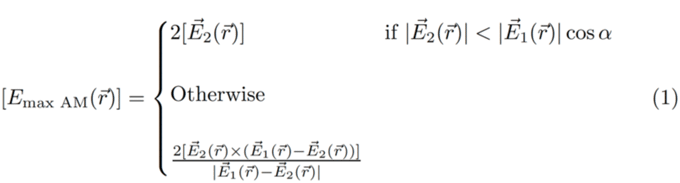

**Equation 1. Amplitude modulation formula.**

### Magnetic Resonance Imaging experiment

#### Participants

A total of 30 healthy volunteers participated in the study, including 18 males (mean = 30.9 ± 9.5 years) and 12 females (mean = 26.2 ± 4.3 years) in accordance with the approved protocol MU-IS/226100/2024/2475492/LF by Masaryk University Research Ethics Committee. Participants completed standard MRI safety screening and health questionnaires to assess contraindications including medication use, neurological/psychiatric history, metal implants, and claustrophobia prior to the MRI session. All participants provided written informed consent prior to the study, in accordance with the Declaration of Helsinki and institutional ethical guidelines.

#### Study design

As illustrated in Figure S5, the study protocol included structural and functional MRI acquisitions, consisting of T1-weighted, T2-weighted, diffusion-weighted imaging (DWI), and simple block MRI or resting-state functional MRI (fMRI). The fMRI protocol was composed of seven runs, each lasting 7 minutes and separated by 3-minute rest intervals. Participants were instructed to remain still, keep their eyes closed, avoid talking or engaging in thoughts related to memory or learning, and not fall asleep. After each run, we assessed participant comfort, verified wakefulness, checked for any discomfort related to electrode placement, and during stimulation runs, we additionally asked whether stimulation was noticed. Participants were encouraged to report any irritation, pressure or heating spots at any time, in which case the session would be interrupted. As shown in the diagram, the first two runs (REST 1 and REST 2) served as a resting-state baseline. After REST 2 and during the subsequent 3-minute interval, stimulation devices were activated, and the amplitude ramped up to reach target intensity by the beginning of STIM 1. The following three runs (STIM 1, STIM 2, and STIM 3) consisted of continuous stimulation periods. After STIM 3, stimulation was ramped down during the next break, followed by two additional post-stimulation runs (POST 1 and POST 2).

#### Experimental setup testing

Before the MRI sessions, we conducted impedance tests using MR-compatible electrodes (Neurocare, 20 mm) and, for comparison, standard Ag/AgCl EEG electrodes (BESDATA, 8 mm). Both types were applied directly to participants’ scalps using TEN20 conductive paste (Weaver and Company) to ensure stable skin contact. Measurements were carried out with a lock-in amplifier (Zurich Instruments MFLI 500 kHz/5 MHz), using a 10 mV signal across frequencies from 1 to 10 kHz. Both electrodes showed similar impedance profiles, decreasing exponentially from approximately 20 kΩ at 1 kHz to around 5 kΩ at 10 kHz. Despite a slightly higher inter-subject variability observed with the MR-compatible electrodes—about 4.6% in the 1–4 kHz range—the average impedance values were comparable: 10.2 kΩ (±5.3 kΩ) for the MR electrodes and 10.5 kΩ (±5.5 kΩ) for the EEG electrodes. We also assessed frequency-dependent current attenuation using a similar setup. A ±1 mA alternating current was applied on the head of a participant through the rubber electrodes used for the MRI experiment, and the resulting current was measured via a differential probe across a 100 Ω shunt resistor and recorded with an oscilloscope. As frequency increased, we observed a consistent reduction in transmitted current: from 2.05 mA at 1 kHz to 1.78 mA at 2 kHz, 1.37 mA at 4 kHz, and down to 0.85 mA at 10 kHz. In case such inconsistency was observed before any TI stimulation in the MRI, the inpout voltage was adapted to match the desired output current. These results highlight the need to verify current delivery across participants, as individual factors, like hair or skin properties can affect conduction, as well as the chosen carrier frequency.

#### MRI acquisition

The acquisition was performed on the Siemens Prisma 3T MR whole-body scanner with 64-channel head-neck coil at Core Facility MAFIL (CEITEC, Masaryk University, Brno, Czechia). High-resolution structural T1 and T2 images were acquired for each participant. Parameters of the T1 MPRAGE sequence were 240 sagittal slices, TR = 2300 ms, TE = 2.34 ms, FOV = 256 mm, flip angle = 8°, voxel size = 0.85 × 0.85 × 0.85 mm. Parameters of the T2 sequence were160 sagittal slices, TR = 3000 ms, TE = 411 ms, FOV = 256 mm and voxel size = 1 × 1 × 1 mm. The BOLD fMRI data was acquired using a multiband T2 echo-planar imaging sequence, with a repetition time of 1126 ms, echo time of 35 ms, voxel size of 2.5 × 2.5 × 2.5 mm, field of view of 208 mm, flip angle of 45 degrees, 60 transverse slices, 370 volumes, and a multiband acceleration factor of 4.

#### Preprocessing and analysis

All MRI data was preprocessed using the Statistical Parametric Mapping 12 (SPM12, The Wellcome Trust Centre for Neuroimaging, London, UK) and Conn toolbox (Whitfield-Gabrieli & Nieto-Castanon, 2012). Anatomical images of each subject underwent an initial visual quality assessment to detect possible artifacts or structural anomalies followed by segmentation, bias correction and spatial normalization to the Montreal Neurological Institute (MNI) space. Functional data preprocessing included realignment (least squares approach and 6 rigid body transformations); correction of susceptibility distortion interactions; outlier detection using ART (0.5 mm framewise displacement threshold); MNI-space normalization; and a 3D Gaussian kernel (5 mm full-width at half maximum, FWHM) smoothing. In addition, functional data were denoised using aCompCor method, including realignment regressors (original, derivative and squared terms), five principal components each from white matter and CSF timeseries, two linear trend terms, and temporal bandpass frequency filtering (0.008 Hz and 0.09 Hz).

First-level analyses were performed using a general linear model (GLM) in SPM for subjects who underwent a block-design stimulation paradigm, consisting of alternating 30-second ON and OFF periods, as well as for those who participated in resting-state scans, that underwent continuous stimulation under different conditions. Each block or resting-state condition was modeled in the design matrix with a canonical hemodynamic response function (HRF) and its temporal derivative (basis set: hrf, order: 16) to account for BOLD response latency variations. Functional connectivity analyses were performed using both seed-based and ROI-to-ROI approaches for 30 subjects randomly assigned to two active stimulation groups (15 participants received TI with a 5 Hz offset and and 15 different participants received TI with a130 Hz offset) across all resting-state sessions. Connectivity estimates were based on Fisher-transformed correlation coefficients obtained from general linear models applied to BOLD signal timeseries, using subject-specific parcellations. Hippocampal regions of interest were obtained from subject-specific segmentations using the FreeSurfer hippocampal subfields and amygdala segmentation tool (Iglesias et al., 2015; Saygin et al., 2017), while cortical regions were defined according to the Harvard-Oxford atlas and HPC-ICA network decomposition. Seed-based connectivity (SBC) was calculated between each seed region and all brain voxels, while ROI-to-ROI connectivity (RRC) was computed between all pairs of ROIs. Group-level analyses employed GLMs with connectivity values as dependent variables and group and condition as predictors. For SBC, statistical inference used Gaussian Random Field theory with a voxel-level threshold of p < 0.001 and cluster-level p-FDR < 0.05. For RRC, connection-level results were considered significant at p-FDR < 0.05.

## 3. Results

### High-frequency stimulation decreases performances in the Rey/Taylor task

The primary outcome measure was the 3-minute recall scores on the Rey/Taylor task assessed either in a baseline (control condition with no stimulation) or assessed during the stimulation trial just after the stimulation was discontinued.

Wilcoxon signed-rank tests were conducted to compare copy and 3-minute recall scores for baseline and inhibitory TI (4-targets / 2-targets hippocampi / 2-targets cortices) conditions. As expected, scores decreased between baseline and copy for all conditions (Tab. 1).

**Table 1:**
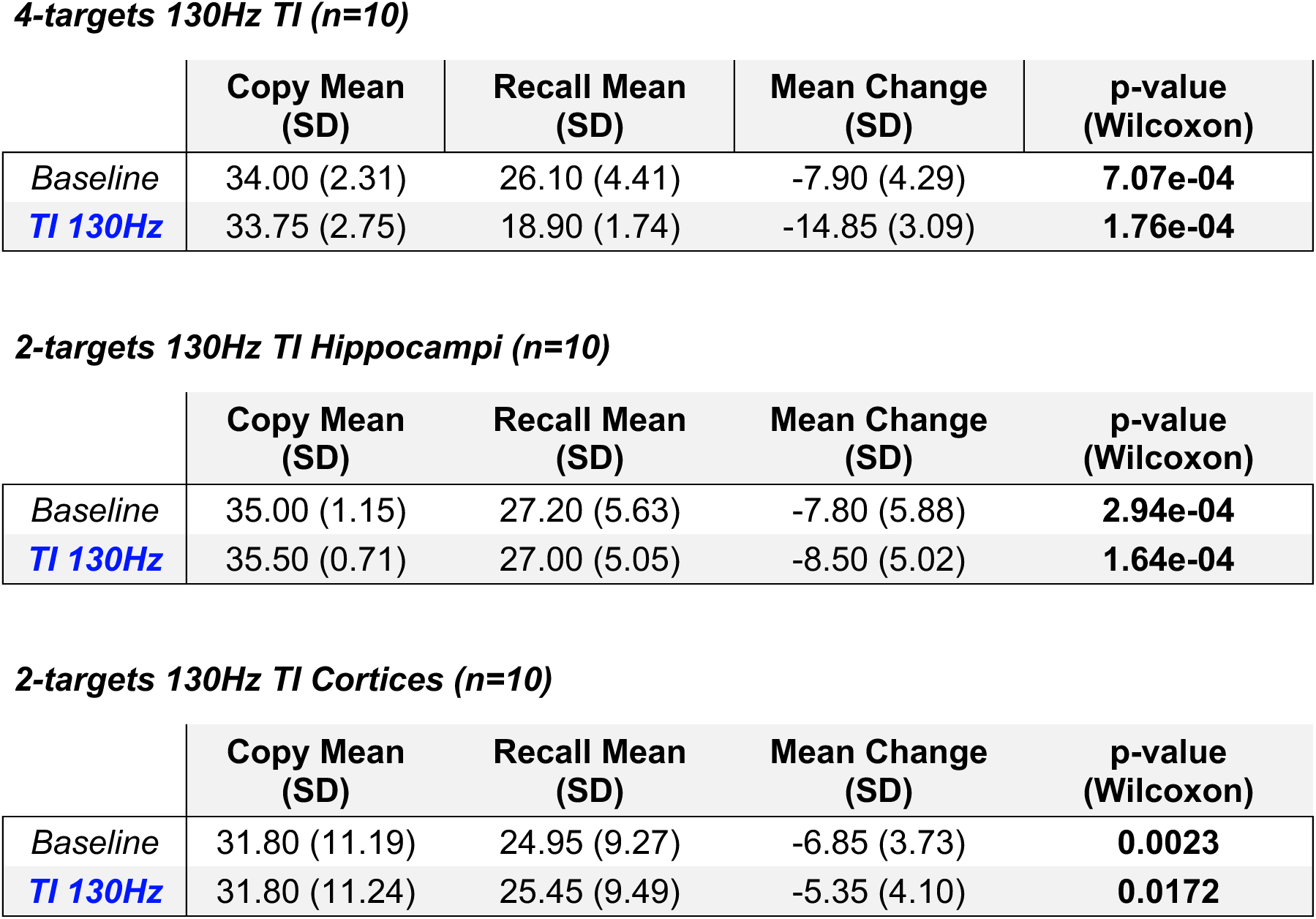
Mean copy and recall scores for the baseline and 130Hz multi-focal TI stimulation.

*4-targets 130Hz TI (n=10)*
|  | Copy Mean (SD) | Recall Mean (SD) | Mean Change (SD) | p-value (Wilcoxon) |
| --- | --- | --- | --- | --- |
| *Baseline* | 34.00 (2.31) | 26.10 (4.41) | -7.90 (4.29) | **7.07e-04** |
| *TI 130Hz* | 33.75 (2.75) | 18.90 (1.74) | -14.85 (3.09) | **1.76e-04** |

*2-targets 130Hz TI Hippocampi (n=10)*
|  | Copy Mean (SD) | Recall Mean (SD) | Mean Change (SD) | p-value (Wilcoxon) |
| --- | --- | --- | --- | --- |
| *Baseline* | 35.00 (1.15) | 27.20 (5.63) | -7.80 (5.88) | **2.94e-04** |
| *TI 130Hz* | 35.50 (0.71) | 27.00 (5.05) | -8.50 (5.02) | **1.64e-04** |

|  | Copy Mean (SD) | Recall Mean (SD) | Mean Change (SD) | p-value (Wilcoxon) |
| --- | --- | --- | --- | --- |
| *Baseline* | 31.80 (11.19) | 24.95 (9.27) | -6.85 (3.73) | **0.0023** |
| *TI 130Hz* | 31.80 (11.24) | 25.45 (9.49) | -5.35 (4.10) | **0.0172** |

**Table 2:**
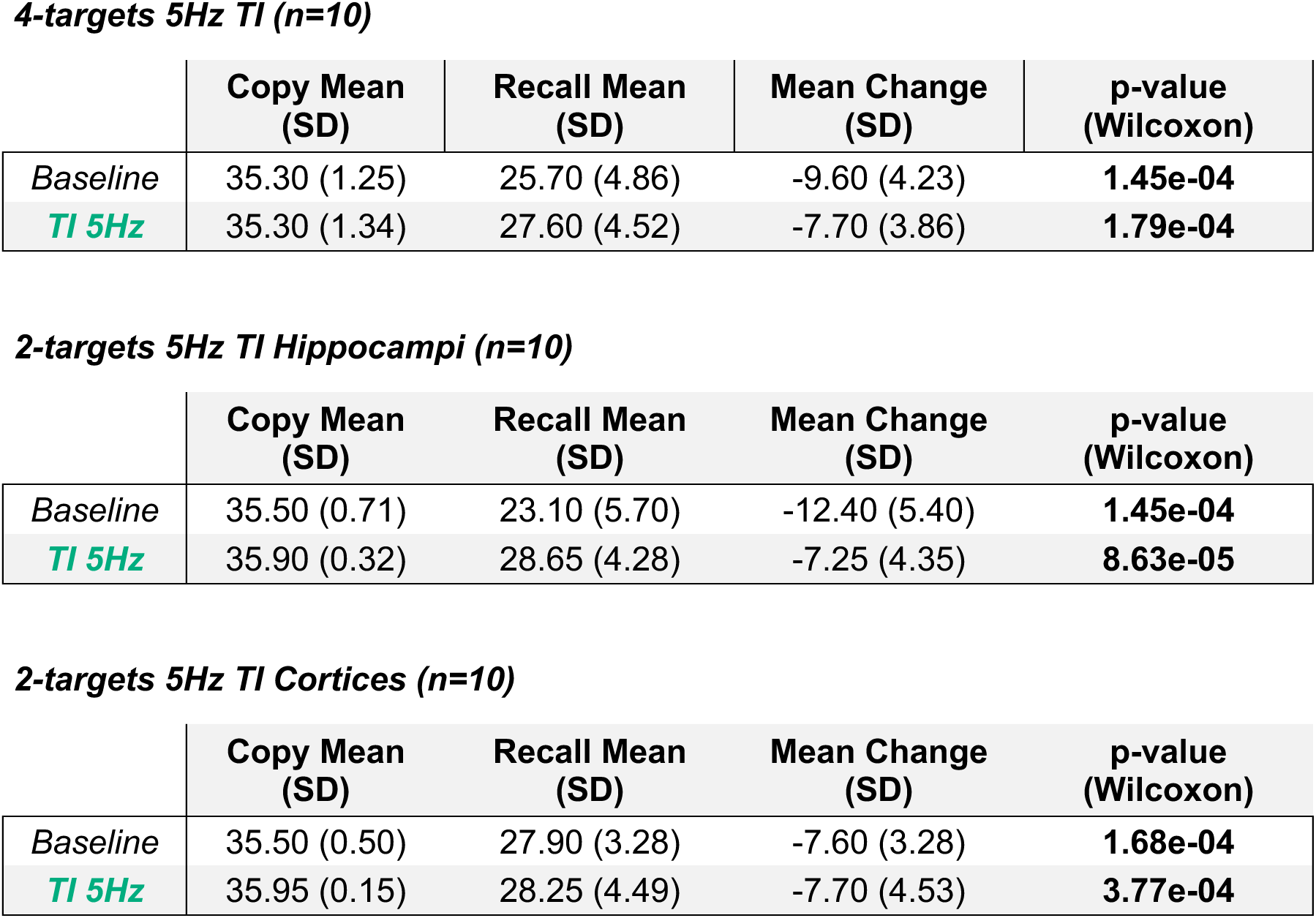
Mean copy and recall scores for the baseline and 5Hz multi-focal TI stimulation.

4-targets 5Hz TI (n=10)
|  | Copy Mean<br>(SD) | Recall Mean<br>(SD) | Mean Change<br>(SD) | p-value<br>(Wilcoxon) |
| --- | --- | --- | --- | --- |
| *Baseline* | 35.30 (1.25) | 25.70 (4.86) | -9.60 (4.23) | **1.45e-04** |
| *TI 5Hz* | 35.30 (1.34) | 27.60 (4.52) | -7.70 (3.86) | **1.79e-04** |

2-targets 5Hz TI Hippocampi (n=10)
|  | Copy Mean<br>(SD) | Recall Mean<br>(SD) | Mean Change<br>(SD) | p-value<br>(Wilcoxon) |
| --- | --- | --- | --- | --- |
| *Baseline* | 35.50 (0.71) | 23.10 (5.70) | -12.40 (5.40) | **1.45e-04** |
| *TI 5Hz* | 35.90 (0.32) | 28.65 (4.28) | -7.25 (4.35) | **8.63e-05** |

|  | Copy Mean<br>(SD) | Recall Mean<br>(SD) | Mean Change<br>(SD) | p-value<br>(Wilcoxon) |
| --- | --- | --- | --- | --- |
| *Baseline* | 35.50 (0.50) | 27.90 (3.28) | -7.60 (3.28) | **1.68e-04** |
| **TI 5Hz** | 35.95 (0.15) | 28.25 (4.49) | -7.70 (4.53) | **3.77e-04** |

Further analysis was conducted to understand the distribution of individual score changes comparing the baseline and stimulation conditions. To compare the magnitude of score changes between the baseline and the 4-target TI condition, we used the a non-parametric ANOVA followed by Wilcoxon rank-sum test. The results showed a significant difference in the ANOVA for session (baseline, TI stimulation), TI protocols (4 targets, 2 target cortices, 2 target hippocampi) and the interaction. Wilcoxon tests later highlight significant differences between the baseline and the 4-target TI condition for 3-minute recall scores (p = 6.54e-04), indicating that the group receiving TI with an offset of 130 Hz in 4 sites simultaneously experienced a significantly greater decrease in performance compared to the baseline condition (without stimulation) (Fig. 2). Using TI with 130 Hz targeting 4 sites (bilateral hippocampus and temporal cortex), 1 participant/10 showed no change (TI vs no stimulation baseline) in recall scores and 9 participants showed a decrease (mean decrease = -7.20, SD = 4.45). Interestingly, none of the 2-targets stimulation groups showed a significant decrease in 3-minutes scores with stimulation (Hippocampi: mean decrease = -0.20 (2.30) – p-value= 0.9394 / Cortices: mean decrease = 0.50 (1.73) – p-value= 0.7033).

No significant correlation was found between age or sex and change in delayed recall scores in any of the conditions. Participants with higher baseline scores showed a more pronounced decrease in performance in the 4-target 130Hz TI condition (r = 0.85), suggesting that initial cognitive performance may modulate the impact of TI stimulation (Fig. 2B).

In summary, the application of multi-focal temporal interference stimulation with 130 Hz offset to the bilateral hippocampi and temporal cortices during encoding resulted in a significant decrease in 3-minute recall scores on the Rey/Taylor task, with the 4-targets stimulation condition experiencing a pronounced reduction in recall ability compared to the baseline condition (Figure 2).

### Low-frequency theta stimulation increases performances in the Rey/Taylor task

TI stimulation with a 5 Hz offset was tested using 4-target or 2-targets protocols like that used for high-frequency stimulation. However, results showed no significant improvement in 3-minute recall (p > 0.05) for the 4-target and bilateral cortex (2-target) groups compared to baseline, indicating that 5 Hz stimulation applied to temporal cortices and hippocampi at the same time or the temporal cortices alone, does not enhance memory performance (Fig. 3B and C).

This outcome is consistent with previous tACS studies indicating that theta stimulation of the temporal cortex can impair memory. For instance, Murray and colleagues (2023) found that theta activity in the left temporal lobe significantly disrupted memory, more than other forms of stimulation. Likewise, Almeida showed that increased theta amplitude in the temporal lobe during recall could interfere with memory processes (Almeida et al., 2023). These findings suggest that theta oscillations are crucial for memory encoding and retrieval and disrupting them in the temporal cortex can negatively affect memory performances.

However, our protocol to focus solely on stimulating the hippocampi showed significant differences, in line with the approach by Violante and colleagues (2023) (Fig. 3D).

Again, across all conditions accuracy followed the expected pattern, with higher scores during copying than recall (p<0.05). As for the high-frequency TI stimulation, a non-parametric ANOVA was realized for delta score ∼ session (baseline, TI stimulation) * TI protocols (4 targets, 2 target cortices, 2 target hippocampus). The ANOVA, as well as the Wilcoxon rank-sum test showed a significant difference in score changes for the session, the protocol and the interaction session*protocol; but more precisely between baseline and bilateral hippocampal stimulation (p = 0.0447), with the bilateral hippocampus TI excitatory stimulation condition exhibiting a significant enhancement in 3-minute recall performance (Fig. 3D).

The individual-level analysis revealed that in the comparison between the baseline and the TI low-frequency conditions, none of the participants showed either a decrease or change in scores, and 10 participants experienced a significant increase (mean increase = 5.55, SD = 2.74). Lower baseline scores were associated with greater improvements in the experimental condition (r = 0.48), suggesting, as for the high-frequency stimulation, that initial cognitive performance impacts stimulation outcome (Fig. 3D).

In conclusion, low-frequency temporal interference stimulation significantly enhanced 3-minute recall performance on the Rey/Taylor task, with the bilateral hippocampal stimulation group demonstrating marked improvements compared to the baseline condition.

#### Comparative Analysis of Stimulation Offset Frequency Effects on 3-minute Recall Scores in the Rey/Taylor Task

In the sham stimulation condition, 10 additional naive participants underwent a stimulation procedure without frequency offset (TI stimulation with Δf=0 Hz). The purpose was to observe any changes in 3-minute recall scores that might occur due to non-specific factors over time and to make sure the high-frequency stimulation (only carrier frequencies) itself does not interfere with memory.

As shown in Figures 1 and S3, sham stimulation induced no significant change in 3-minute recall scores (p = 0.5687), indicating that the observed effects in the experimental conditions are unlikely to be attributed to non-specific factors such as participant fatigue or increased task familiarity. This specificity underscores the bidirectional potential of TI stimulation, which can either enhance or disrupt memory performance based on stimulation parameters. Differences in 3-minute recall from the sham condition were compared to both experimental conditions, revealing significant enhancements with excitatory stimulation (p5Hz vs sham = 4.89e-04) and significant disruptions with inhibitory stimulation (p130Hz vs sham = 0.0027). Furthermore, the 3-minute recall scores for each stimulation offset were compared through a Wilcoxon rank-sum test (Table 3), which confirmed a significant difference in score changes between the excitatory hippocampi TI (Δf=5 Hz) and inhibitory 4-targets TI (Δf=130 Hz) conditions (p = 1.81e-04), highlighting how stimulation frequency critically determines whether TI boosts or impairs memory encoding.

**Table 3:** Mean immediate 3-minute recall scores for the different stimulation frequencies offset (130Hz, 5Hz, and 0Hz).

|  | Baseline RM<br>(SD) | Stimulation RM<br>(SD) | Mean Change<br>(SD) | p-value<br>(Wilcoxon) |
| --- | --- | --- | --- | --- |
| **5Hz 2-targets**<br>**hippocampi TI**<br>**(n=10)** | 35.90 (0.32) | 28.65 (4.28) | -7.25 (4.35) | **8.63e-05** |
| **130Hz 4-targets**<br>**TI (n=10)** | 33.75 (2.75) | 18.90 (1.74) | -14.85 (3.09) | **1.76e-04** |
| **0Hz 4-targets TI**<br>**(n=10)** | 28.20 (4.73) | 27.65 (4.86) | -0.55 (2.67) | 0.5687 |

RM = Recall Mean

Additionally, low-frequency (Δf=5 Hz) TI stimulation targeting only the bilateral hippocampus (2 targets) during the behavioral experiments significantly enhanced 3-minute recall performance on the Rey/Taylor task, whereas high-frequency (Δf=130 Hz) stimulation targeting both the bilateral hippocampus and temporal cortex (4 targets) produced a significant decrease in scores. This opposing pattern demonstrates the bidirectional modulation achievable with TI: enhancement through focused, low-frequency activation of key memory hubs versus disruption via broader, high-frequency interference across memory networks. The sham stimulation data further support the specificity of these bidirectional effects, confirming that the observed improvements or impairments are not due to non-specific influences. These findings emphasize the pivotal role of stimulation frequency and target focality in achieving desired cognitive outcomes, positioning TI as a versatile tool for both memory enhancement and targeted disruption in clinical and research applications.

Finally, we examined the relationship between baseline recall scores and the change in recall scores following stimulation to further illustrate bidirectionality at the individual level. For the inhibitory stimulation (130 Hz, 4-target), there was a strong negative correlation between baseline scores and the change in recall scores (r = -0.85, p < 0.001), indicating that participants with higher baseline performance experienced greater disruptions in recall after stimulation. In contrast, for the excitatory stimulation (5 Hz, 2-target hippocampal), there was a moderate positive correlation (r = 0.48, p < 0.01), suggesting that participants with lower baseline scores benefited more from the enhancement. This differential correlation pattern reinforces how TI can amplify strengths in one direction while exacerbating vulnerabilities in the other, tailored by frequency and targeting. Moreover, these findings are consistent with prior work with electrical stimulation with implanted depth electrodes that suggests the baseline performance of the patient will determine whether a desired effect can be achieved (e.g., enhancing memory effects have been reported to occur only in patients whose baseline scores were impaired) (Inman et al., 2018).

The fMRI data provide critical insights into the neural mechanisms underlying the behavioral effects of TI stimulation on memory encoding. As depicted in Figure 4, excitatory TI stimulation targeting the bilateral hippocampi with a 5 Hz offset frequency (carrier frequencies: left 4000/4005 Hz, right 2000/2005 Hz; amplitude ±2 mA) resulted in a significant increase in Blood-Oxygen-Level-Dependent (BOLD) signal within these regions (p<0.05, non-parametric Friedman ANOVA, corrected). The fMRI experiment, conducted using a block design (30s ON/30s OFF), revealed robust activation in the medial temporal lobe, particularly in the hippocampi, as evidenced by axial brain slices and the corresponding BOLD time series extracted from these areas. In a standard block-design fMRI GLM (stimulation blocks interleaved with rest), the hippocampi shows a positive effect for the contrast Stim (5 Hz) > Rest (T=1.64, p<0.005), i.e., higher BOLD during stimulation relative to the interleaved rest baseline (example shown for one participant in Figure 4). This heightened hippocampal activity aligns with the behavioral findings, where participants exhibited significantly improved recall scores on the Rey-Taylor figure task following excitatory TI stimulation compared to sham.

**Figure 4:**
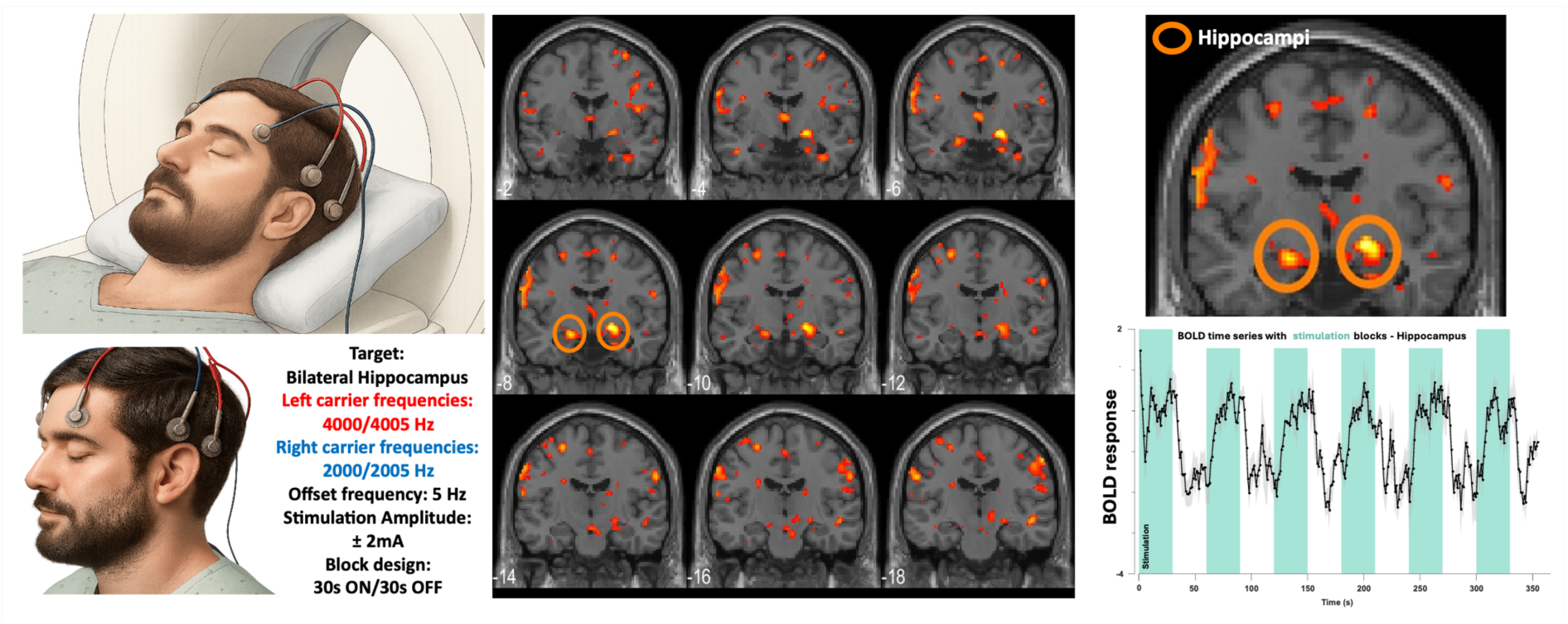
Induced changes in BOLD signal and Functional Brain connectivity following Temporal Interference stimulation of bilateral hippocampi with a 5 Hz offset. Temporal interference stimulation can be adapted to fMRI environment to modulate specific brain regions and/or brain-wide activity. TI has been used to target bilateral hippocampi (one carrier frequency per hippocampus) to modulate hippocampal memory networks. In a standard fMRI block design where the stimulation is provided in blocks, bilateral hippocampi show a significant increase in BOLD activity compared to baseline – highlighting the capability of TI stimulation to induce region-specific activity changes. In a separate Resting-State protocol, TI stimulation of the same bilateral hippocampi could induce whole brain changes with wide changes of functional connectivity within the hippocampal, default mode and attentional network. *Human images in this figure are computer-generated and do not represent any participants in the study*.

Additionally, resting-state fMRI indicated increased functional connectivity within the hippocampal, default mode, and attentional networks following stimulation, suggesting that TI not only activates localized brain regions but also modulates large-scale networks involved in memory and attention.

To further elucidate the neural mechanisms underlying the behavioral effects of TI stimulation on memory encoding, we analyzed functional connectivity changes with resting-states fMRI sessions during 5 Hz (excitatory) and 130 Hz (inhibitory) stimulation compared to a resting state baseline. As shown in Figure 5 and S7, both frequencies induced significant connectivity alterations, with distinct memory network patterns (Yeo et al., 2011).

**Figure 5:**
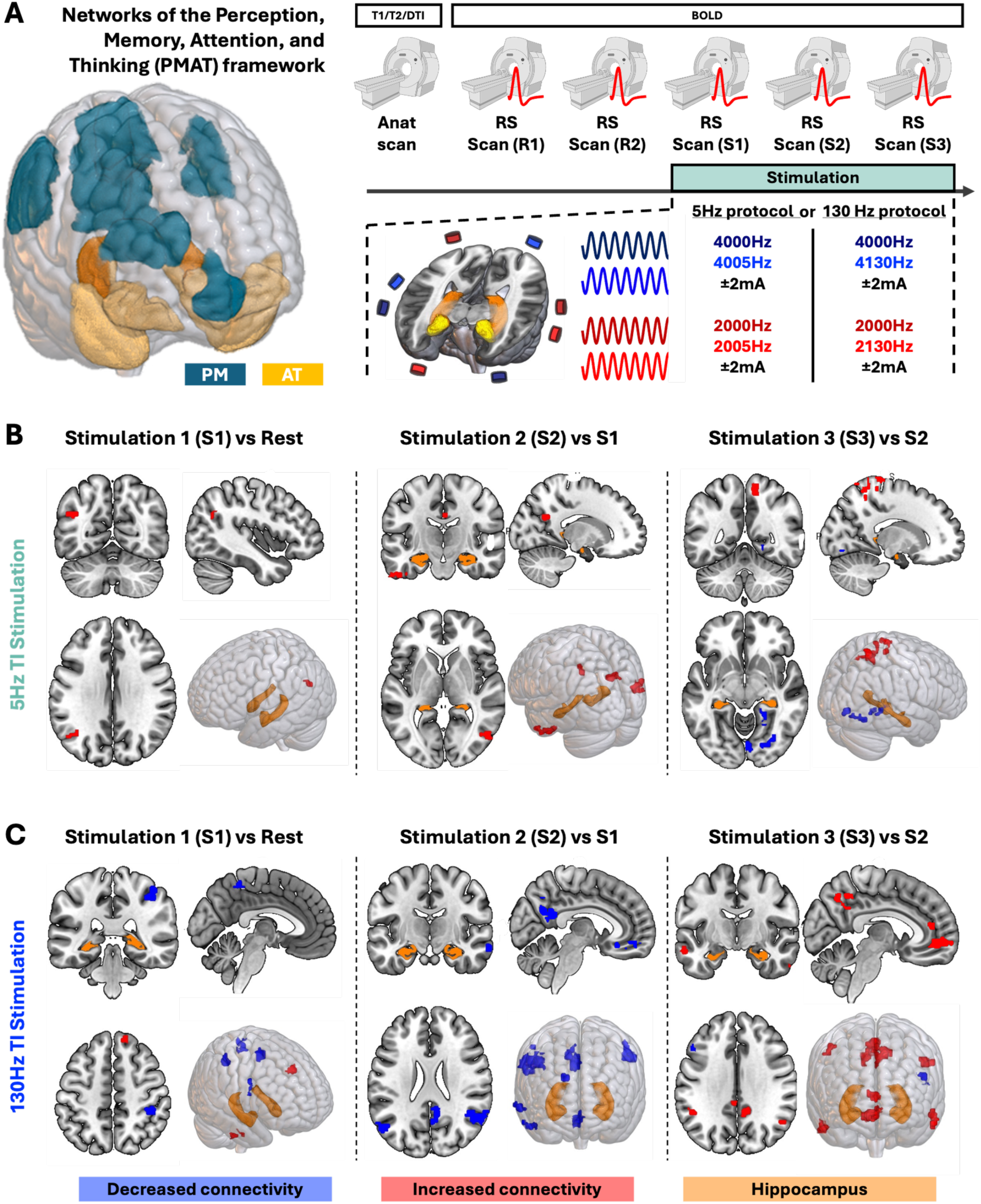
Multi-focal Temporal Interference Stimulation Effects on Brain Connectivity. Multi-Focal Temporal Interference stimulation can modify brain connectivity, derived from Resting-State (RS) fMRI data of 30 healthy participants (n=15 per stimulation frequency group). The connectivity was calculated using the Head Body Tail hippocampi parcellated ROIs (Iglesias et al., 2015; Saygin et al., 2017). Each hippocampus was used as a seed to evaluate the connectivity changes between all the other ROIs. The maps are representing both hippocampi activations, i.e., right and left (with their subdivisions). Bilateral hippocampi were targeted with high frequency or low frequency TI, and the effect of the stimulation was assessed within the PMAT networks (Inhoff & Ranganath, 2017), highlighting subregions in panel A left (PM in cyan, AT in yellow) (**A**). Functional connectivity changes (FDR corrected, p-value<0.05) during 5 Hz (excitatory) and 130 Hz (inhibitory) stimulation, with blue denoting decreased connectivity, red indicating increased connectivity, and orange marking the hippocampus are displayed in B and C (Rest 1 and Rest 2 have been concatenated into Rest). Connectivity shifts are highlighted with a global increased connectivity within the PMAT networks for 5 Hz stimulation and a decreased FC for the 130 Hz stimulation. *R1: Rest 1/ R2: Rest 2/ S1: Stimulation 1/ S2: Stimulation 2/ S3: Stimulation 3*.

For 5 Hz stimulation, which enhanced memory recall, we observed robust connectivity increases within and between memory-related networks of the PMAT (Figure 5).

For Stim 1 vs. Rest, the right hippocampus (seed) exhibited increased resting-state connectivity with the occipital cortex and angular gyrus (Δ ≈ 0.11 z-Fisher; t(14)=5.44, p<0.001; d≈1.1–1.4), consistent in 13/15 participants (≈87%) and representing ∼68% of the Rest mean. For Stim 2 vs. Stim 1, the left hippocampus showed increased connectivity with the left inferior temporal gyrus (Δ ≈ 0.12 z-Fisher; t(14)=8.38; p<10⁻⁶; d≈1.9–2.2) in all 15 participants, and with left pITG, aITG, aMTG, and pMTG (Δ ≈ 0.14 z-Fisher; t(14)=8.28; p<10⁻⁶; d≈1.7–2.1) in 14/15 participants; and the precuneus cluster also increased (Δ ≈ 0.13 z; t(14)=6.65; p<0.001; d≈1.16–1.72) in 14/15 participants. Using the left hippocampus head as seed, connectivity rose to right iLOC and sLOC (Δ ≈ 0.14 z; t(14)=5.92; p<0.001; d≈1.5–1.6) in 14/15 participants. The right hippocampus body increased with the posterior cingulate gyrus (Δ ≈ 0.12 z-Fisher; t(14)=5.57; p<0.001; d≈0.80–1.44) in 14/15 participants, while the left pITG and pTFusC cluster rose (Δ ≈ 0.12 z; t(14)=8.43; p<10⁻⁶; d≈2.1) in all 15 participants. For Stim 3 vs. Stim 2, the left hippocampal tail displayed decreases in three posterior clusters (lingual gyrus, right occipital fusiform gyrus, right hippocampus; Δ ≈ −0.11 z; |d|≈1.4–2.1, but an increase in a superior cluster (precuneus, right superior parietal lobule, lateral occipital cortex; Δ ≈ +0.10 z; t(14)=9.97; d≈2.6) in all 15 participants. The right hippocampal tail showed increases in three clusters (right superior parietal lobule, postcentral gyrus, lateral occipital cortex, precentral gyrus; Δ ≈ +0.12–0.13 z; p≈3×10⁻⁵; d≈1.4–1.6) in 14/15 participants.

In addition, we observed changes in connectivity in broader networks like the Default Mode Network (DMN) – encompassing the hippocampus, medial prefrontal cortex, and posterior cingulate cortex – and the Fronto Parietal Network, with effects persisting after FDR correction; these results are consistent with literature of network activity changed during enhanced memory states (Dresler et al., 2017)(Figure S7).

Conversely, 130 Hz stimulation, did not induce FC change when comparing the gathered stimulation sessions to the baseline resting state. However, when splitting the stimulation sessions, we observed primarily altered connectivity in the PMAT networks and sensory/motor networks, including regions such as the Superior Temporal Gyrus, Heschl’s Gyrus, and Precentral Gyrus in the first 7 minutes of stimulation (S1). Further sub-analyses of 130 Hz stimulation revealed dynamic connectivity shifts across different stimulation conditions.

For Stim 1 vs. Rest, the left hippocampus exhibited stronger connectivity with the right frontal pole (+0.13 z, p=0.0001), right inferior temporal gyrus (+0.11 z, p≈6.9e-06), and right paracingulate gyrus (+0.12 z, p≈1.9e-06), but weaker connectivity with the right postcentral/supramarginal gyrus (−0.10 to −0.12 z, p≈7.3e-05 to 1.0e-05), right precentral/inferior frontal gyrus (−0.13 z, p≈2.2e-05), right middle/superior frontal gyrus (−0.13 z, p≈5.1e-06), left precuneous/postcentral gyrus (−0.11 z, p≈1.1e-06), and left lateral occipital/superior parietal lobule (−0.11 z, p=0.0001). For Stim 2 vs. Stim 1, the left hippocampus showed reduced connectivity with the right angular/lateral occipital gyrus (−0.15 z, p≈2.9e-06), right frontal pole (−0.12 z, p=0.0001), right middle temporal gyrus/temporal pole (−0.14 z, p≈1.7e-05), left lateral occipital cortex (−0.13 z, p≈2.9e-07), cingulate gyrus/precuneous (−0.13 to −0.16 z, p≈9.9e-05 to 2.2e-05), and subcallosal/paracingulate gyrus (−0.15 z, p≈6.3e-05). The right hippocampus had diminished connectivity with the frontal medial/paracingulate gyrus (−0.14 z, p=0.0001) but enhanced connectivity with the right precentral gyrus (+0.15 z, p=6e-06), left superior temporal gyrus (+0.12 z, p=9e-06), right lateral occipital/superior parietal lobule (+0.12 z, p=3.7e-05), left planum polare/temporal pole (+0.12 z, p=3.4e-05), and right inferior frontal gyrus (+0.15 z, p=7e-05). For Stim 3 vs. Stim 2, the left hippocampus demonstrated stronger connectivity with the precuneous/cingulate gyrus (+0.14 z, p≈3.7e-06), right middle temporal gyrus/temporal pole (+0.12 z, p≈1.8e-06), left supramarginal/angular gyrus (+0.11 z, p≈3.7e-05), left middle temporal gyrus/temporal pole (+0.11 z, p≈1.5e-05), and right angular/lateral occipital gyrus (+0.11 z, p≈1.3e-04), but weaker connectivity with the left middle/inferior frontal gyrus (−0.12 z, p≈4.7e-06). The right hippocampus showed enhanced connectivity with the cingulate/paracingulate gyrus (+0.17 z, p≈5.5e-05), right frontal pole/paracingulate gyrus (+0.17 z, p<1e-06), precuneous/cingulate gyrus (+0.14 z, p<1e-06), right lateral occipital/angular gyrus (+0.11 z, p≈1.5e-05), right paracingulate gyrus (+0.13 z, p≈4.8e-05), frontal medial/paracingulate gyrus (+0.12 z, p≈2.9e-05), subcallosal/paracingulate gyrus (+0.11 z, p≈3.6e-05), and left frontal orbital/insular cortex (+0.12 z, p≈4.8e-05).

A potential explanation of the above results could be that the 130 Hz stimulation induces a brain-wide decrease in functional connectivity that gets compensated during the stimulation protocol. Within the PMAT networks, temporal and mesial regions show an increased FC during S3 (Figure 5) – this phenomenon also recurs brain-wide with the cortical regions showing a broad increase in FC (Figure S7).

These findings suggest that TI stimulation modulates brain networks in a frequency-dependent manner, with excitatory 5Hz stimulation enhancing memory network connectivity to support improved encoding, while inhibitory 130Hz stimulation shifts connectivity toward non-memory networks, potentially disrupting memory processes when applied brain-wide.

## 4. Discussion

This study demonstrates frequency- and target-dependent, bidirectional modulation of figural memory by non-invasive multi-focal temporal interference (TI). During encoding, 5 Hz envelope stimulation targeted to the bilateral hippocampi enhanced 3-minute Rey/Taylor recall, consistent with theta-range facilitation of hippocampal-cortical communication (Suthana et al., 2012; Jun et al., 2019; Liu et al., 2012; Jun et al., 2020; Ezzyat et al., 2018; Ezzyat et al., 2017). In contrast, 130 Hz envelope stimulation delivered concurrently to four sites (bilateral hippocampi and anterior temporal cortices) impaired recall, whereas two-target hippocampal stimulation at 130 Hz—similar to envelope frequencies used therapeutically in Parkinson’s disease and epilepsy—did not affect performance. Baseline-dependent effects were observed: inhibitory 130 Hz showed a strong negative correlation with baseline ability (r = −0.85), while excitatory 5 Hz showed a moderate positive correlation (r = 0.48), suggesting susceptibility to disruption at higher baseline and greater benefit at lower baseline, respectively (Cleary et al., 2025).

Converging fMRI results clarify mechanisms linking frequency and network engagement. Periodic 5 Hz hippocampal TI tracked BOLD variation locally and increased resting-state connectivity within hippocampal, PMAT, default-mode (DMN), and attentional networks, aligning with theta’s role in synchronizing encoding-relevant assemblies (Rugg & Vilberg, 2013). By contrast, 130 Hz preferentially shifted connectivity toward PMAT and sensory/motor systems (including superior temporal gyrus, Heschl’s gyrus, and precentral gyrus), with within-session dynamics (Stim2 vs. Stim3) suggestive of compensatory reweighting outside core memory circuits; these network shifts mirror the observed behavioral impairment. The pattern is consistent with frequency-specific modulation of encoding and/or early consolidation: high-frequency TI can down-regulate ripple-range processes linked to memory formation (Acerbo et al., 2022; Missey & Acerbo et al., 2024), potentially perturbing cortico-hippocampal interactions essential for both encoding and later retrieval (Goyal et al., 2017; Jacobs et al., 2016; Wang et al., 2015; Khodagholy et al., 2017; Missey et al., 2025; Kunz, 2024). These findings extend prior human TI work showing focal 5 Hz hippocampal modulation with memory benefits (Violante et al., 2023).

Together, the results establish that TI can bidirectionally “tune” memory networks—enhancing encoding with focal 5 Hz hippocampal stimulation while disrupting performance with distributed 130 Hz multi-target stimulation—and motivate state- and trait-informed personalization of parameters. Beyond basic science applications that dissect anatomical and physiological substrates of memory (including subjective metamemory phenomena; Neisser et al., 2022), clinical avenues include targeted enhancement of memory in health and disease and, conversely, disruption of maladaptive memory encoding (e.g., post-traumatic stress), provided parameters are precisely timed and individualized (Laxton et al., 2013; Widge et al., 2019). More broadly, the frequency-by-focality principle we delineate—focal, on-target theta for enhancement without deficit; multi-site high-frequency for inhibition—offers a compact framework for rational, non-invasive deep neuromodulation design.

### Limitations of the Study

While our study provides valuable insights into the effects of Temporal Interference stimulation on memory encoding, several limitations should be considered. The study was conducted in healthy participants, constraining generalizability to clinical populations. The recall delay was short (3 min), limiting inferences about long-term consolidation and clinical durability of effects. Inter-individual variability in electrode placement and neuroanatomy may have introduced heterogeneity in current distributions, a particular concern for multifocal TI, and could affect reproducibility across cohorts and sites although consistent results have been shown over the 100 participants in the present study.

MRI-specific constraints required different carrier frequencies while holding the offsets constant (5 Hz for excitatory, 130 Hz for inhibitory), to ensure safe operation and avoid interference or heating; prior work indicates that neuromodulatory effects primarily track the offset rather than the carrier, supporting this approach (Missey et al., 2025). Hardware also limited us to bilateral hippocampal targeting in MRI (rear head-coil port incompatible with 16-lead four-target configurations). Future studies should use individualized, MRI-guided montage optimization and modeling to reduce anatomical variance, extend retention intervals to probe consolidation, include clinical cohorts, and enable true four-target stimulation in-scanner to better assess therapeutic potential and mechanistic specificity (Missey et al., 2025).

## 5. Conclusion

Multi-focal TI can bidirectionally modulate human memory in a manner governed by both frequency offset and anatomical specificity. Focal 5 Hz (theta-range) stimulation of the bilateral hippocampi enhances performance, consistent with the role of hippocampal theta in organizing unit activity and coordinating cortico-hippocampal communication during encoding. In contrast, distributed 130 Hz stimulation across hippocampi and temporal cortices disrupts recall, indicating that high-frequency offsets can perturb network dynamics subserving mnemonic processing. Together, these findings articulate a frequency-by-focality principle: low-frequency, on-target hippocampal stimulation supports encoding without measurable deficit, whereas multi-site high-frequency stimulation impairs performance.

This controllable, bidirectional influence on memory provides a mechanistic foothold for both basic and translational work. In cognitive neuroscience, TI affords causal tests of circuit- and network-level hypotheses about encoding and retrieval. Clinically, it motivates parameterized interventions that either bolster memory function or selectively dampen maladaptive encoding, contingent on individualized anatomy and brain state. The approach thus outlines a rational path toward non-invasive, state-informed neuromodulation for memory enhancement and for targeted disruption where therapeutically indicated.

## 6. Acknowledgments

This work was supported by the Ministry of Health of the Czech Republic in cooperation with the Czech Health Research Council in the context of the RECOVERY project under No. NW25J-04-00094 (F.M.). A.W. received funding from the European Union’s Horizon Europe research and innovation programme under grant agreements No. 101088623 (EMUNITI), No. 101157945 (FITSLEEP), and from the Ministry of Health of the Czech Republic in cooperation with the Czech Health Research Council in the context of the SENSE project under No. NW25-08-00053. Additionally, this work was supported by the Specific University Research Grant, as provided by the Ministry of Education, Youth and Sports of the Czech Republic in the year 2025 (MUNI/A/1683/2024). Finally, this work received financial support from Agence Nationale de la Recherche under France 2030 bearing the reference “ANR-24-RRII-0005”, on funds administered by Inserm. N.P.P. was supported by NIH K08NS105929. D.L.D. was supported by NIH R01 NS088748. I.R.V was supported by the BBSRC (BB/Y011856/1).

## 7. Authors Contribution

F.M. and A.W. conceived and designed the project. F.M., E.J-M., C.L., J.T., M.A.S., A.A., O.S., S.K. conducted human experiments. F.M., E.J-M., and A.A. analyzed data. F.M. created the finite-element models. F.M. and E.J-M. wrote the first draft of the manuscript and refined it after inputs from C.L., J.T., M.A.S., A.A., O.S., S.K., I.R.V., M.B., V.J., G.T., D.L.D, N.P.P., and A.W.

## 9. Data Availability Statement

All data produced in the present study are available upon request to the authors.

## 11. Supplementary Figures

**Figure S1:**
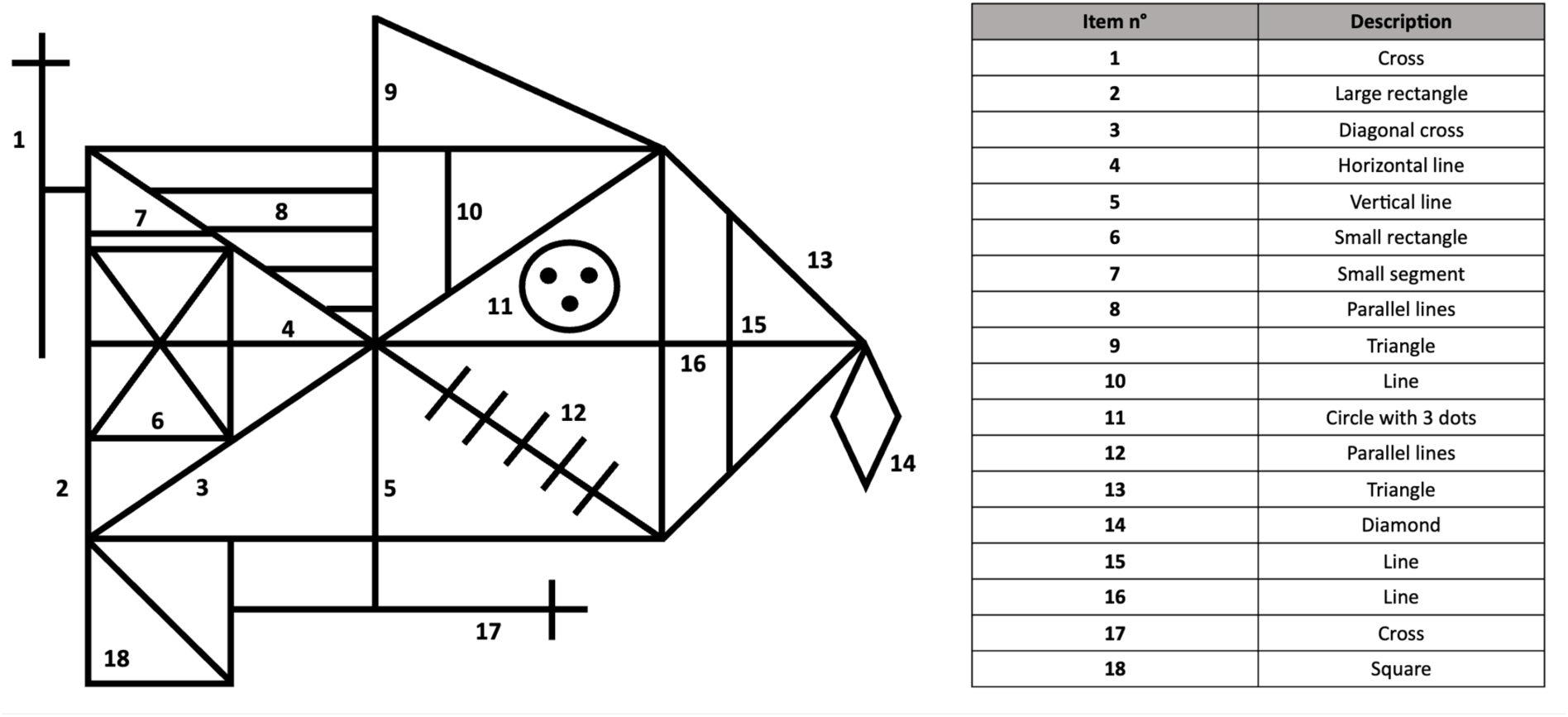
Rey Figure scoring rules (Osterrieth, 1944).

**Figure S2:**
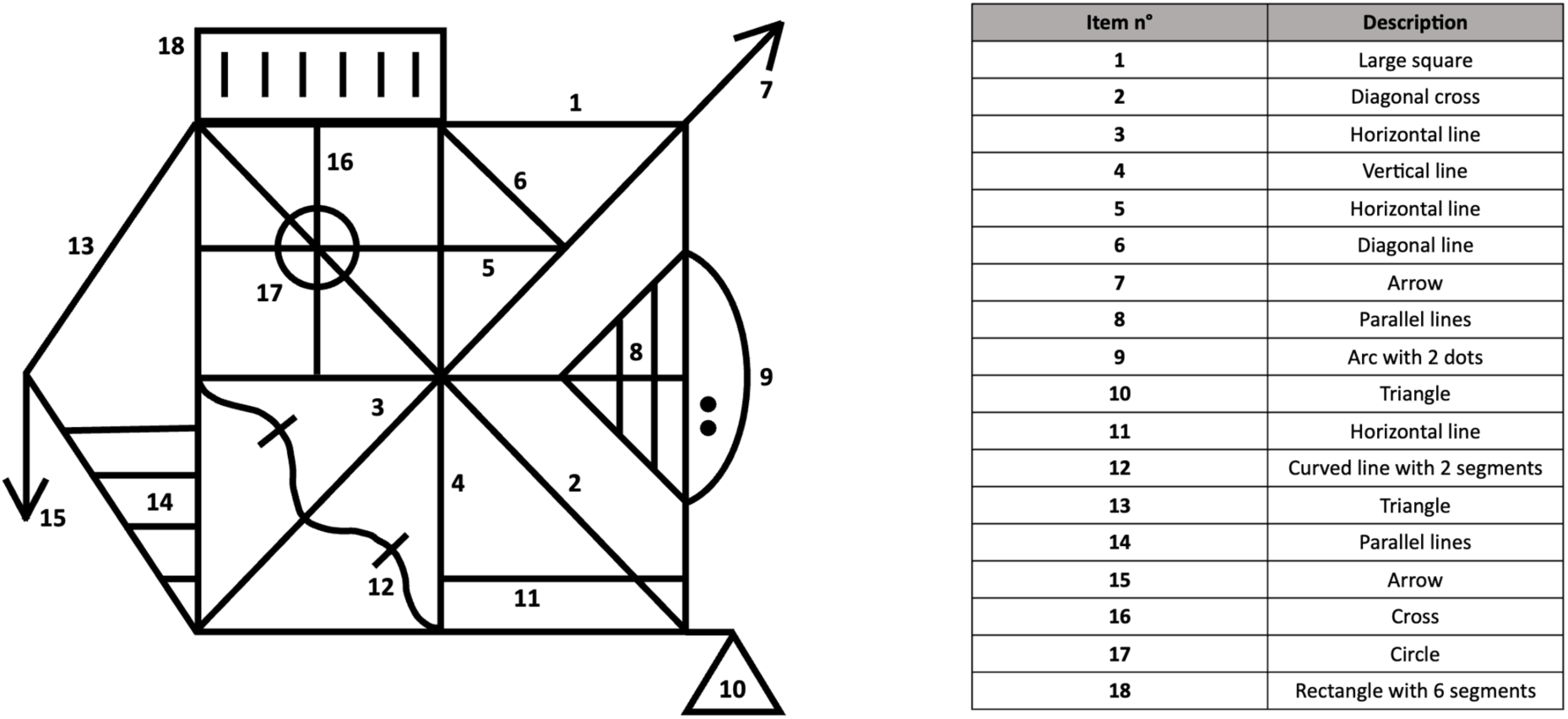
Taylor Figure scoring rules (Taylor, 1969).

**Figure S3:**
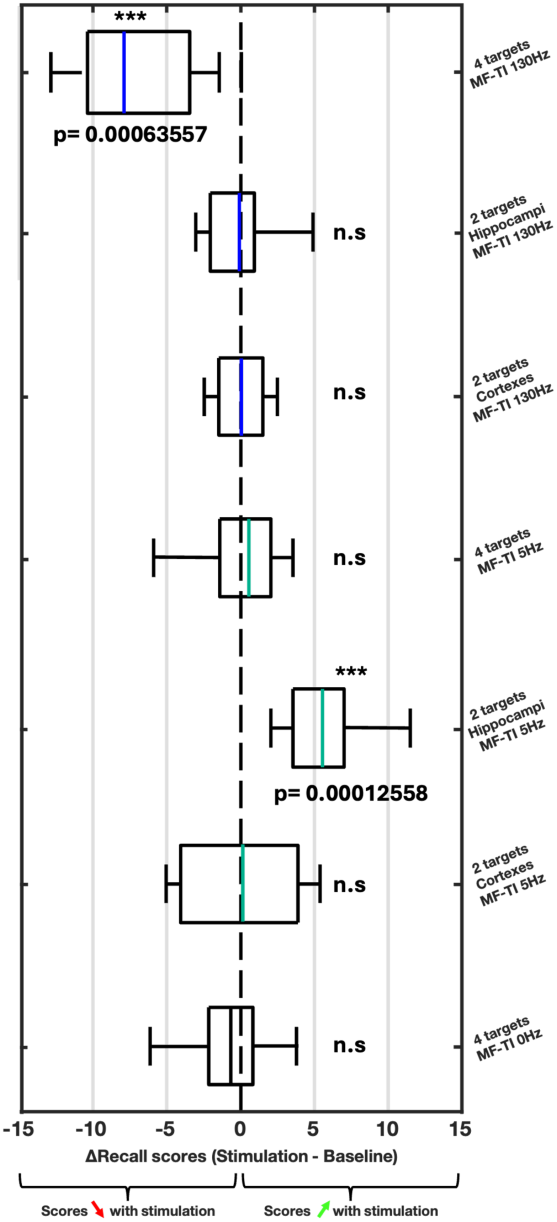
Forest plot of the frequency-dependent effect of the multi-focal TI stimulation on Rey/Taylor figure scores. Boxplots showing differences in recall scores between baseline and stimulation sessions for all protocols (high-frequency 130 Hz, low-frequency 5 Hz, sham). Left shifts indicate reduced performance (e.g., high-frequency four-target), right shifts indicate improved performance (e.g., low-frequency hippocampal), with sham showing no effect.

**Figure S4:**
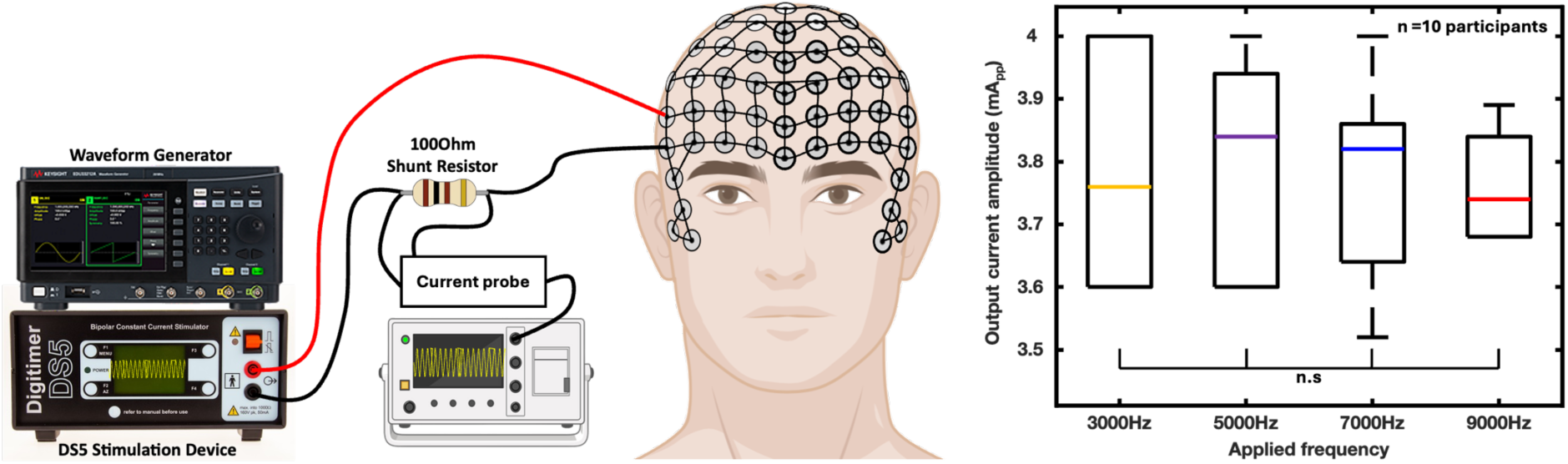
Recording of the output current across stimulation electrodes for different carrier frequencies. Boxplots of output current measured across a 100Ohm resistor for carrier frequencies of 3kHz, 5kHz, 7kHz, and 9kHz over 10 participants, with intended output set at 4mApp and actual currents varying within a 0 to -13% error margin, confirming consistency in stimulation delivery.

**Figure S5:**
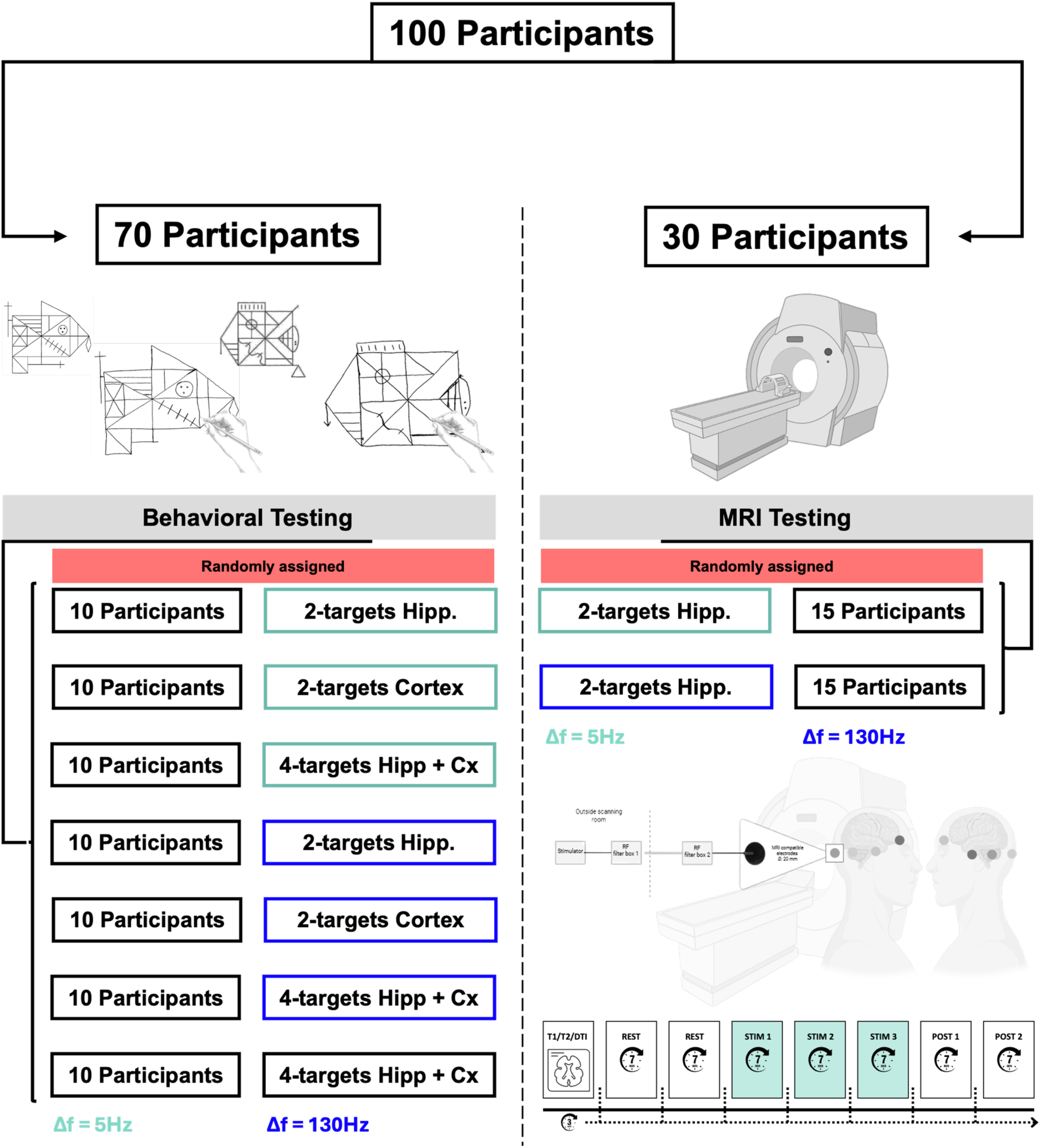
Experimental design and randomization of the study. 100 participants were divided into two groups, one of 70 participants for the behavioral study and one of 30 participants for the MRI study. Within the two different studies, participants were again randomly assigned to either of the stimulation protocol groups (7 groups of 10 participants or 2 groups of 15 participants). The MRI study design is also depicted in the figure, with anatomical MRI followed by 7 resting-state scanning runs – in which 3 are set during the TI stimulation session. Pale green references to 5Hz offset stimulation and blue references to 130Hz offset stimulation.

**Figure S6:**
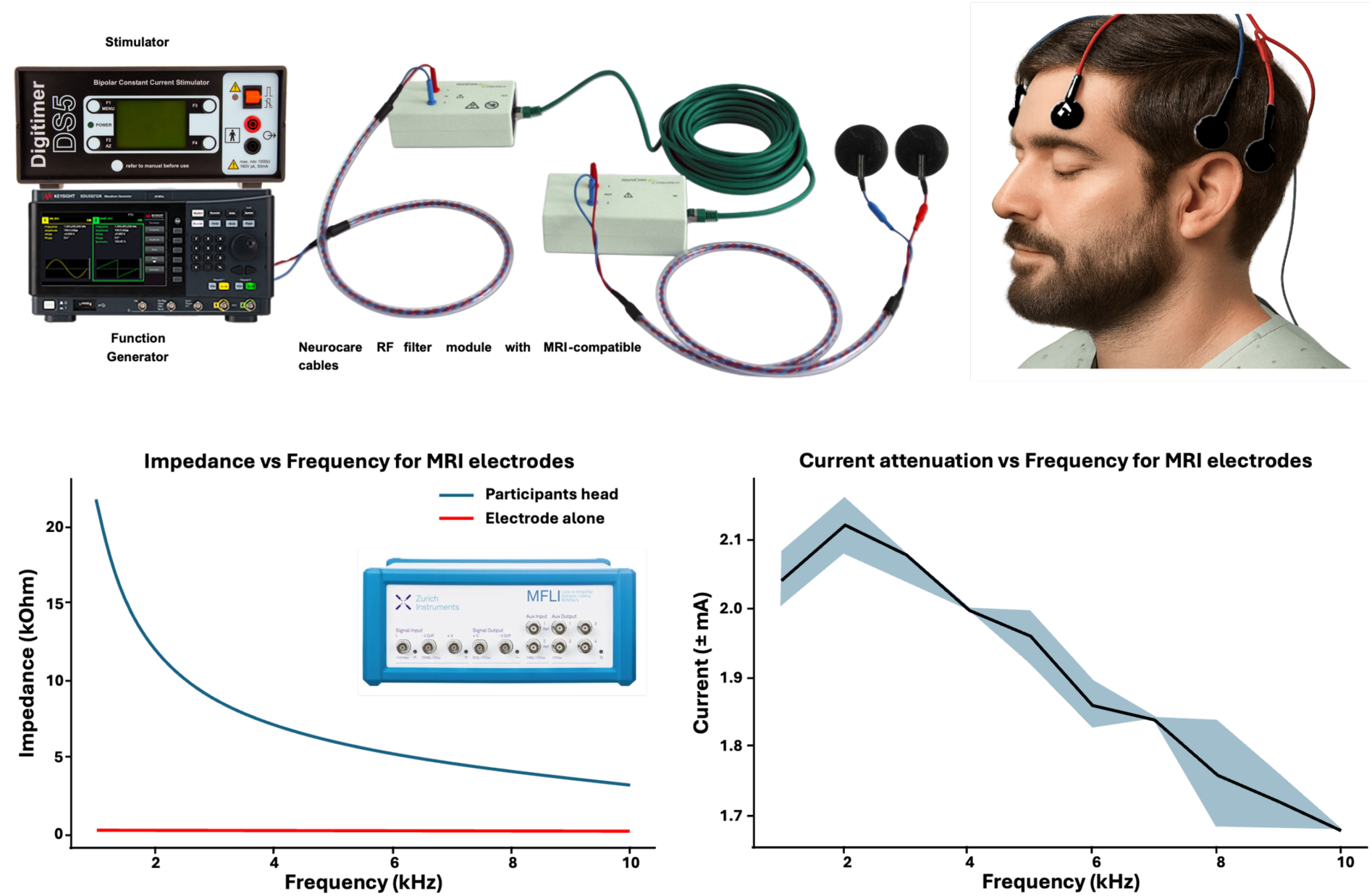
MRI compatible stimulation electrode characterization. MRI compatible electrodes were characterized in the fMRI environment to assess the impedance of the stimulation system and the current attenuation of the stimulation. Based on the recorded attenuation, stimulation frequencies were chosen to better match the necessary current output. *Human images in this figure are computer-generated and do not represent any participants in the study*.

**Figure S7:**
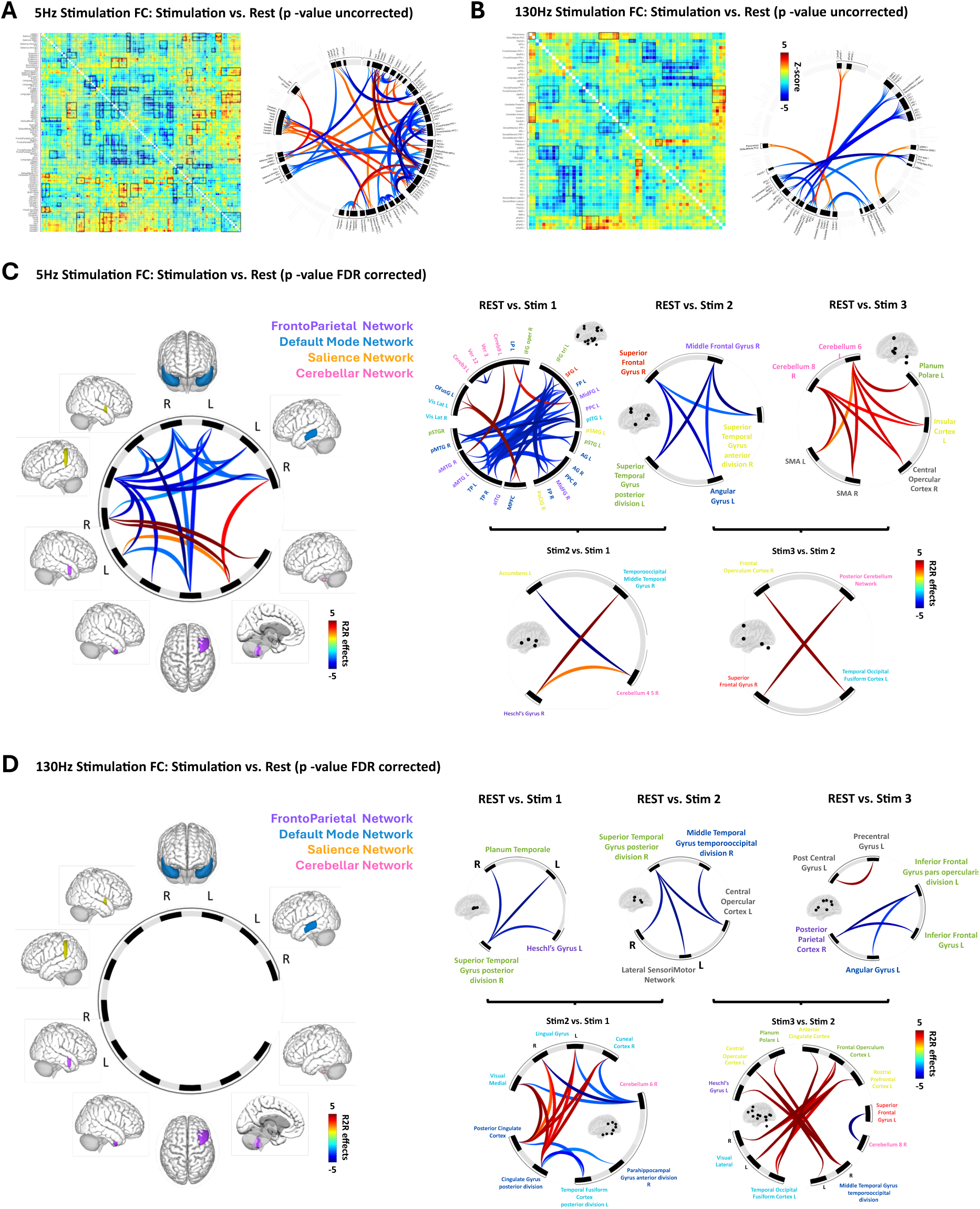
Functional Connectivity Changes Induced by 5Hz and 130Hz Multi-Focal Temporal Interference Stimulation. Functional connectivity alterations during Multi-Focal Temporal Interference stimulation compared to baseline resting state, assessed using fMRI with a lower effect for the stimulation at 130Hz (**A** and **B**). Connectivity matrices and circular plots for 5Hz stimulation (excitatory) show enhanced connectivity within the Default Mode Network (DMN), including the hippocampus, medial prefrontal cortex, and posterior cingulate cortex, and the FrontoParietal Network, with significant changes surviving FDR correction (**C**). Matrices and plots for 130Hz stimulation (inhibitory) highlight altered connectivity in sensory and motor networks, including the Superior Temporal Gyrus, Heschl’s Gyrus, and Precentral Gyrus, with dynamic shifts noted in sub-analyses (e.g., REST vs. Stim1, Stim2 vs. Stim1). Connectivity strength is represented by Z-scores (-5 to 5), with red indicating positive and blue indicating negative correlations (**D**). These patterns provide a neural basis for the observed memory enhancement with 5Hz and impairment with 130Hz stimulation.

